# Impact on *S. aureus* and *E. coli* membranes of treatment with chlorhexidine and alcohol solutions: insights from molecular simulations and nuclear magnetic resonance

**DOI:** 10.1101/2022.08.31.505867

**Authors:** Callum Waller, Jan Marzinek, Eilish McBurnie, Peter J Bond, Philip T. F. Williamson, Syma Khalid

## Abstract

Membranes form the first line of defence of bacteria against potentially harmful molecules in the surrounding environment. Understanding the protective properties of these membranes represents an important step towards development of targeted anti-bacterial agents such as sanitizers. Use of propanol, isopropanol and chlorhexidine can significantly decrease the threat imposed by bacteria in the face of growing anti-bacterial resistance *via* mechanisms that include membrane disruption. Here we have employed molecular dynamics simulations and nuclear magnetic resonance to explore the impact of chlorhexidine and alcohol on the *S. aureus* cell membrane, as well as the *E. coli* inner and outer membranes. We identify how sanitizer components partition into these bacterial membranes, and show that chlorhexidine is instrumental in this process.

## INTRODUCTION

Pathogenic bacteria pose an enormous threat to human, animal, and plant health, especially given the rate at which resistance to current antibiotics is developing.[1,2] Effective methods of sanitization, such as antimicrobial hand and body scrubs, are essential in the control of widespread diseases as they offer a route to decreasing the rates of infection through human contact.[3–5] To develop new and more effective sanitizing agents, it is essential to understand their mechanisms of action against different bacterial membranes.

Chlorhexidine (CHX) is a member of a chemically related group of antimicrobials called bisbiguanides that form a di-cation at physiological pH (Figure 1).[6] The structure of CHX consists of two chlorophenyl (CPL) functional groups, each bonded to a separate biguanide (BGU) connected *via* a hexane (HEX) linker. Its structure and cationic charge are thought to be key to its bactericidal properties.[7] CHX is widely used as an antiseptic and disinfectant due to its broad efficacy, affordability, and safety, and it is commonly found in mouth washes and surgical scrubs.[8,9] It is effective against both Gram-positive and Gram-negative bacteria. CHX is thought to achieve bacterial cell death by binding to the lipid bilayer head groups and disrupting the membrane, leading to leakage of cellular contents. Hence, it is often combined with alcohol which is theorised to work in a similar way.[10] Under physiological pH the two BGU groups carry a cationic charge of 1 *e* each. The cationic nature of the molecules is proposed to play a major role in their interaction with negatively charged bacterial membranes.[11]

**Figure 1.**
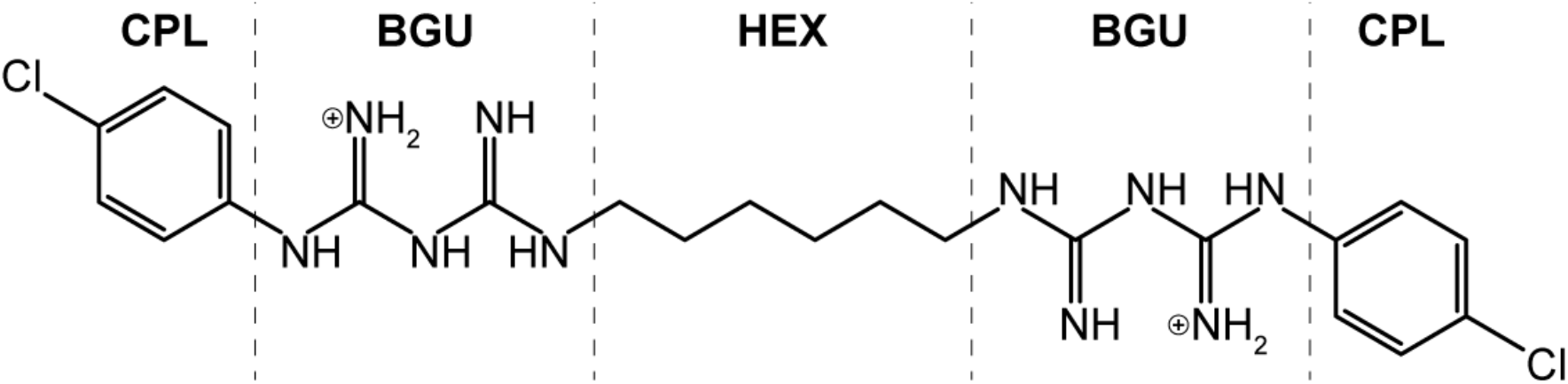
Separation of CHX into key functional groups which are necessary in the interaction of CHX with a PL membrane.

Short chain alcohol molecules are theorised to kill bacteria primarily by disrupting the membrane, similarly to CHX.[12] The value of topically applied alcohol as an antibacterial agent is well known, and has been proven as an effective form of defence against a plethora of pathogens *in vivo*, including *E. coli* and *S. aureus*.[3–5,13] This is largely attributed to macroscale perturbations which are imposed upon the bacterial cytoplasmic membrane, a model which was expanded upon by Feller *et al*.[14] who tracked the permeation of alcohol across a POPC membrane *via* nuclear Overhauser enhancement (NOE) spectroscopy.[15] This showed that alcohol accumulates mostly around the phosphate region of the phospholipid (PL) membrane, with the carbon chains of alcohol orientated toward the membrane centre. Further to this, it has been shown that the introduction of short-chained alcohols to a PL membrane can significantly affect the surface area per lipid (APL) and membrane thickness.[16] More recently this phenomenon was supported using molecular dynamics (MD) simulations by Ghorbani *et al*.[17] This also showed that following diffusion from the bulk ethanol solution there is very little diffusion of alcohol back out of the membrane plane.

CHX is theorised to bind at the headgroup region of membranes *via* the CPL and BGU functional groups as a ‘wedge’, parting the headgroups and creating gaps which make the membrane ‘leaky’.[18] This charge interaction between the BGU region and the bilayer was shown by Oosten *et al*.[19] to result in binding further from the membrane centre but more strongly to the head group region in more negatively charged membranes. It has been demonstrated *via* cell culture experiments and verified by Franz diffusion cell studies that unlike other antiseptics, CHX can remain active due to epidermis binding even after the bulk of the sanitizer solution has been washed away.[20,21] CHX is active against *E. coli* (archetypal Gram-negative bacteria) and *S. aureus* (archetypal Gram-positive bacteria).[11] These two examples present three different kinds of membranes with alternative lipid compositions to CHX; the inner (EcIM) and outer membrane (EcOM) of *E. coli*, and the cell membrane of *S. aureus* (SaCM). CHX in sanitizing agents is available in both alcohol and aqueous solutions.

The EcOM contains lipopolysaccharide (LPS) in the outer leaflet and PLs in the inner leaflet, whereas the EcIM and SaCM contain PLs in both leaflets. The PLs in both the inner leaflet of the EcOM and both leaflets of the EcIM are phosphoethanolamine (PE), phosphoglycerol (PG) and cardiolipin (DPG) in a ratio of 90:5:5.[22] The precise details of the LPS chemical structure are dependent upon the *E. coli* strain. Briefly, the polar region of LPS is composed of layers of sugars, some of which are phosphorylated, whereas the hydrophobic portion typically has six hydrocarbon tails.[23,24] The SaCM is composed of PG, lysyl-phosphatidylglycerol (LPG) and DPG in a ratio of 54:36:10.[25,26] The main difference between the SaCM and the EcIM is the presence of LPG instead of PE, which provides the capacity for head group hydrogen bonding between LPG and PG while also lowering the overall membrane charge.[27]

To gain molecular insight into the mechanism(s) of action of CHX on the membranes of *E. coli* and *S. aureus*, here we have employed a combination of atomistic MD simulations and solid-state nuclear magnetic resonance (NMR) to study CHX in aqueous and alcohol solutions with the three membrane types presented by the two bacterial species. We present data detailing the partitioning of alcohol and chlorhexidine into bacterial membranes *via* a multidisciplinary approach, using MD and NMR in tandem to determine both their conformational and orientational properties. Our data helps to rationalize the origins of the characteristic membrane deformations generally associated with the use of alcohol and CHX as sanitising agents.

## RESULTS

### Interaction between chlorhexidine and the bacterial membranes in aqueous solution

200 ns simulations were performed in triplicate with CHX in 0.15 M KCl was applied on either side of the three membranes (SaCM, EcIM and EcOM). Visual inspection of the system after 200 ns revealed little perturbation of the membranes in any of the simulations. The insertion depth was evaluated by calculating the CHX density in the z dimension (parallel to the membrane normal). Figure 2(a) shows greater penetration into the sub-head group region of the SaCM compared to either of the *E. coli* membranes. There was a shift in the CHX density towards the centre of the membrane as the simulation proceeds, while no such shift was observed for the *E. coli* membranes. The deeper penetration of CHX may be due to the higher negative charge in the headgroup region of the SaCM (−0.3 e per lipid), compared to the *E. coli* PL leaflets (−0.15 e per lipid) enabling stronger stabilising electrostatic interactions. This is in agreement with previously reported findings by Van Oosten *et al*.[19] that CHX binds more strongly with increased charge disparity. There was no penetration into the sub-headgroup region of the outer leaflet of the OM, despite the highly charged nature of LPS. The slow-moving, almost impenetrable nature of LPS has been well-documented in both experimental and simulation studies.[28] For the PL membranes, over the course of 200 ns the CHX molecules transitioned from partial interaction with the membrane on one terminal BGU-CPL region, to mostly being intercalated and bound by both termini, as illustrated in Figure 2(b). In this configuration, the BGU and CPL functional groups are buried in/beneath the phosphate region toward the ester functional groups, while the HEX group lies along the bilayer surface. The interaction observed here has been termed as a ‘c-shape’ mode of binding and is reminiscent of the wedge proposed by Komljenović *et al*.[18] It should be noted, however, that the wedge they proposed is an inverted conformation, with the CPL-BGU region located in the lipid headgroups and the HEX functional group buried in the lipid tail region.

**Figure 2.**
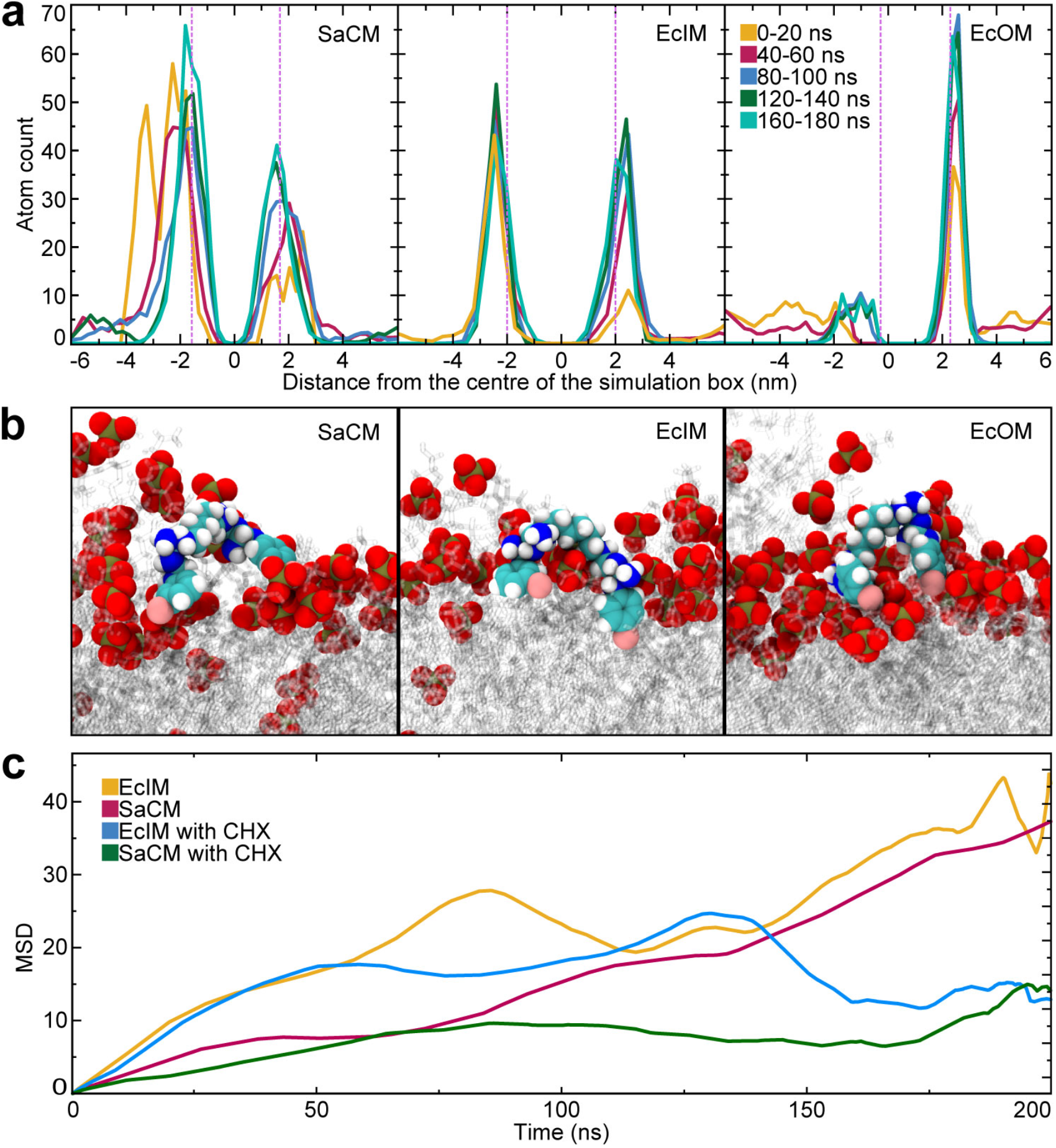
Analysis of the SaCM, EcIM and EcOM when exposed to CHX in a 0.15 M solution of KCl. (A) 20 ns block average density of CHX at 40 ns intervals along the z-axis of the simulation box for the entire 200 ns production run with average head group phosphate position as a pink, dotted line. (B) Snapshots of CHX C-shape binding in the (B1) SaCM, (B2), EcIM and (B3) EcOM systems are shown as spheres with phosphate oxygen in red, phosphate phosphorous in gold, carbon in cyan, nitrogen in blue, chlorine in pink, hydrogen in white and lipids as translucent.

The solvent accessible surface area (SASA) of the CHX was calculated for the simulated trajectories to assess any potential clustering of CHX molecules, but this revealed no net change upon membrane interaction, indicating that CHX aggregation was not induced by their interaction with lipids. Some CHX-CHX interactions between the BGU and CPL regions were observed, but these did not persist for longer than a few nanoseconds (see Supplementary Figure 1). The density of the membrane components as a function of distance from the membrane centre was compared at the start and end of the simulations, revealing no obvious change (see Supplementary Figure 2). Thus, binding of CHX to the membranes studied here did not cause any real displacement of the membrane components. Nevertheless, calculation of the mean squared displacement (MSD) of lipids in the EcIM and SaCM revealed that CHX significantly diminished lipid diffusion suggesting that its lipid binding reduces the mobility of lipids in the bilayer due to the c-shape binding. This confirms an effect which has been suggested from previous research.[29] It should be noted that this decrease was also significantly greater in the SaCM, which was seen to experience CPL binding at greater depth.

The average APL of each membrane was calculated. The SaCM and EcIM membranes only exhibited a marginal difference when CHX was added which is further elaborated upon in sections pertaining to these membranes (see Supplementary Figure 3 and Figure 4). The EcOM did not show any significant difference between systems with or without CHX.

### Chlorhexidine in alcohol solutions: Action on the *S. aureus* cell membrane

Sanitizer solutions which contain CHX are often found as alcoholic formulations. Given the intrinsic antibacterial activity of alcohol,[30–32] it is of interest to determine the additional impact of CHX. We therefore performed 200 ns comparative simulations of the three bacterial membranes in the presence of 20% w/v propanol (PROH) and isopropanol (ISOP) solutions with and without CHX, in triplicate.

Simulation systems containing the SaCM were surrounded by 0.5% w/v CHX in 0.15 M KCl with either 20% PROH or ISOP in bulk water on either side of the membrane. Visual inspection revealed bilayer distortion occurring almost immediately (t = 16.5 and 42 ns for PROH and ISOP, respectively), and becoming more apparent throughout the 200 ns simulation. This phenomenon was also clear when measuring the APL (see Supplementary Figure 3 and Figure 4). Over the course of simulations, the lipids reoriented in a manner that disrupted the canonical bilayer structure of the membrane. This was evident from a gradual increase in APL which plateaued at a value approximately 80% greater than the equilibrated membrane had at t = 0 ns for both PL membranes. It should be noted that due to the hydrogen bonding between PG and LPG in the SaCM, the initial deformation rate was slower; these hydrogen bonds hold the headgroups together and provide an initial resistance to the dispersion caused by alcohol. Alcohol molecules became distributed in the lipid core region of the now very distorted bilayer, allowing the CPL regions of CHX to bind beneath the membrane surface at the headgroup-tail interface, as shown in Figure 2(b). This binding occurred regardless but was more rapid and significant when alcohol was present. The membrane deformation was characterized quantitatively by plotting the membrane density along the simulation cell z-axis (parallel to the membrane normal, see Figure 3(a)). System density plots also showed that alcohol distributed within the membrane by adopting energetically favourable positions at the headgroup/tail interface where it could maximise its energetically favourable interactions (Figure 3 (b)). This positioning of PROH allowed the alcohol functional group to remain in the phosphate/ester region and maintain polar interactions, while its hydrocarbon tail was able to form lipophilic interactions with the lipid tails region, minimising interactions with water. To illustrate this, the lipids in the SaCM model were mapped by their contacts with PROH and water. This clearly shows that alcohol accumulated significantly more around the headgroup/tail interface, where we propose it to take energetically stable conformations (Figure 4(a-c)). Snapshots of the proposed hydrogen bonding occurring during simulations can be seen in Supplementary Figure 5. A similar effect was observed for systems containing ISOP, although the change was less dramatic since the branched structure of secondary alcohols does not align as effectively with the lipid tails (see Supplementary Figure 6). In Figure 4(d) the location of PROH partitioning in the membrane was determined by plotting the minimum distance of a random PROH molecule with POPG headgroups, sub-headgroups and tail regions. This shows that while alcohol partitioned within the bilayer (regions highlighted in cyan) it is in closest contact with the sub-headgroup and tail regions (as defined in Figure 4(d)) of the lipids. This was performed for 5 PROH molecules in total which can be seen in Supplementary Figure 7. These results agree with similar computational work performed by Ghorbani *et al*.[17] which found ethanol to accumulate at the headgroup/tail interface.

**Figure 3.**
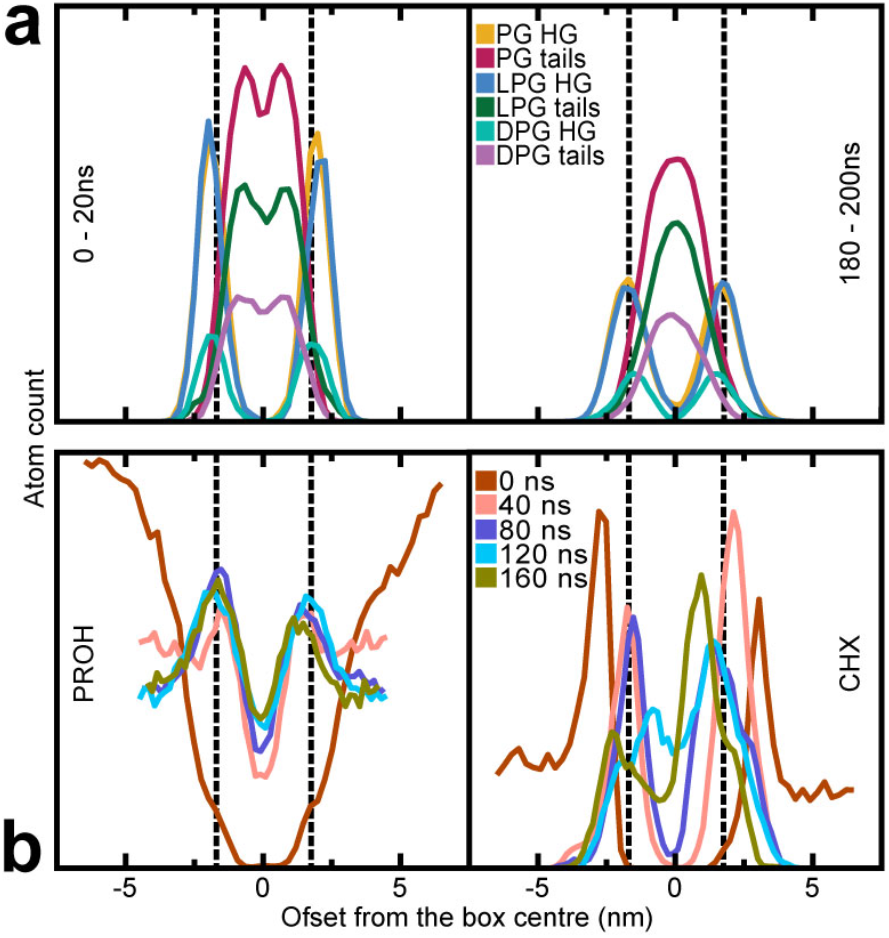
Analysis of the density along the z-axis of the SaCM system exposed to separate solutions of PROH and CHX in 0.15 M KCl. (A) Density of membrane components along the z-axis of the simulation box of at 0 and 180 ns averaged over the following 20 ns. (B) The density of PROH and CHX at 40 ns intervals in production, averaged over the following 20 ns.

**Figure 4.**
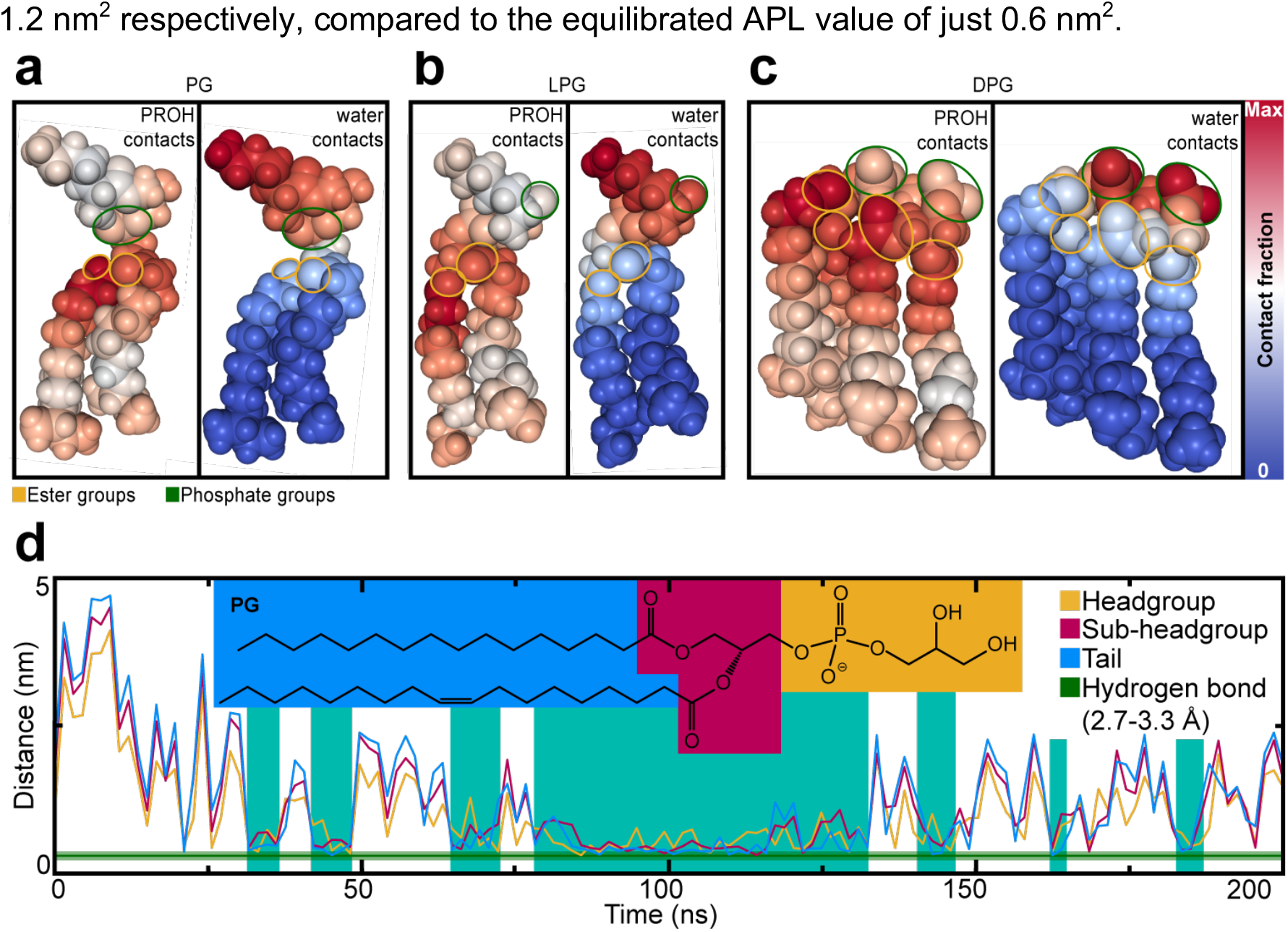
Analysis of the SaCM when acted upon by separate solutions of PROH and CHX in 0.15 M KCl. (a-c) Contact mapping of (a) PG, (b) LPG and (c) DPG in the SaCM bilayer between 100 and 200 ns with PROH and water mapped linearly from (blue) 0 contacts to (red) the most seen by any atom in the lipid, showing ester groups circled in yellow and phosphate groups circled in green. (d) Distance from a random PROH molecule to the (yellow) headgroup, (red) interface and (blue) tail regions of PG in the bilayer, tracked over the full 200 ns production run and indicating the hydrogen bonding distance (2.7-3.3 Å)[67] in green and periods of extended interaction in cyan.

Analysis of system densities showed that CHX aggregated around the bilayer headgroups (Figure 3 (b)) where it interacted with the SaCM *via* the same c-shape interactions observed in the absence of alcohol. Over the course of the 200 ns production run CHX molecules became bound by both of their BGU/CPL moieties which then became embedded within the headgroup-tail interface of the membrane (see Supplementary Figure 8). Interestingly, CHX ‘bridging’ was observed; i.e. PROH caused significant deformation and thinning of the membrane such that CHX molecules bound to separate leaflets were able to contact each other across the bilayer, interacting *via* their BGU/CPL regions. However, there appeared to be no additional deformation compared to the alcohol systems which did not contain CHX. This is reminiscent of the ‘handshake’ binding mode proposed by Komljenović *et al*.[18] Their findings suggest that CHX could operate *via* a similar mechanism to alcohol, by penetrating the membrane and aggregating in the membrane centre. However, this was not seen in simulations of CHX without alcohol. The SASA was again tracked to determine whether CHX molecules had any tendency to cluster, but no suggestion was seen other than a dip corresponding to a binding cluster of 4 CHX molecules which dissipated within 30 ns of forming on the bilayer surface (see Supplementary Figure 9). This was not seen in repeats but suggested the absence of clusters was due to CHX forming interactions with the membrane in preference to those it would form with itself. The APL was also examined (Figure 5(a)), revealing a consistent, gradual increase in APL in both PROH and ISOP systems, resulting in a plateau at 1 and 1.2 nm^2^ respectively, compared to the equilibrated APL value of just 0.6 nm^2^.

**Figure 5.**
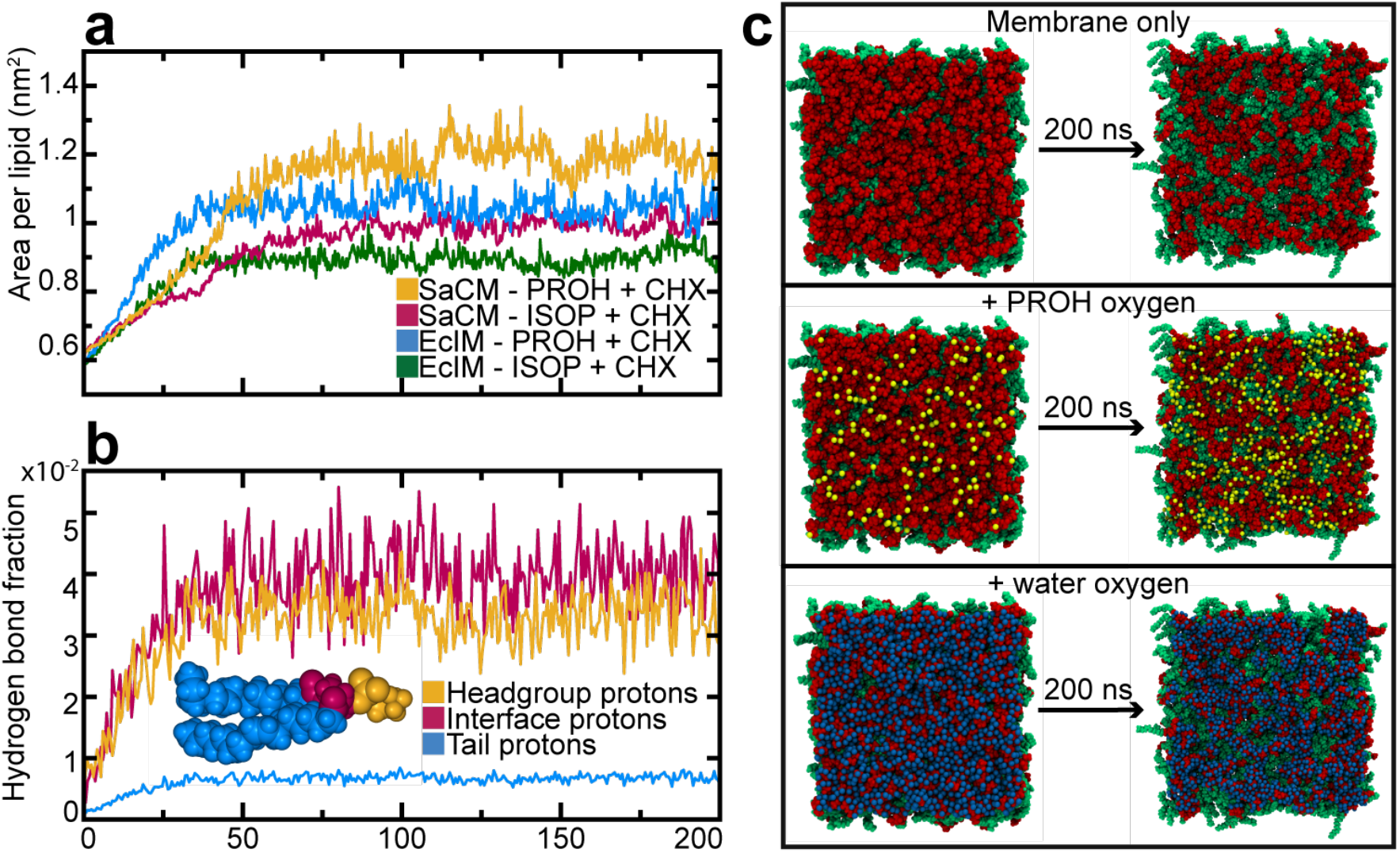
(a) Average APL of lipids in the SaCM and EcIM when exposed to CHX in (yellow and blue respectively) PROH and ISOP over the full 200 ns production run. (b) Hydrogen bonds from PROH to POPE in the EcIM divided by the number of protons in that region to standardise to a per proton basis, showing the tails in blue, the interface in red and the headgroup in yellow. (c) Visual snapshots of the EcIM surface exposed to CHX in 20% PROH at 0 ns and 200 ns with all solution hidden, showing PROH oxygen and showing water oxygen, showing headgroups in red, lipid tails in green, PROH oxygens in yellow and water oxygens in blue.

### Chlorhexidine in alcohol solutions: action on the *E. coli* inner membrane

The same approach was employed for the EcIM (i.e. membranes simulated in the presence of 0.5% w/v CHX in 0.15 M KCl and either 20% PROH or ISOP). A reduced final APL was observed as the simulations progressed, but this was a faster and more substantial initial increase compared to the lipids of the SaCM (Figure 5(a)), likely because of the lower membrane charge of the EcIM (−0.15 e per lipid) compared to the SaCM (−0.3 e. Thus, the EcIM was impacted less by the alcohol/alcohol+CHX compared to the SaCM (see Supplementary Figure 10 and Figure 11). There was also a slightly slower decrease in the average SaCM Scc order parameter, again pertaining to LPG-PG hydrogen bonding resisting initial deformation. As with the SaCM membrane, alcohol was located at the headgroup/tail interface in the EcIM (Figure 4(c-e), Supplementary Figure 12). This was also evident from measurement of the system density which showed less aggregation of CHX on the membrane surface of the EcIM compared to the SaCM, again presumably due to the lower membrane charge (see Supplementary Figure 13 and Figure 14). The same partitioning conformation of alcohol as seen in the SaCM system was observed again (see Supplementary Figure 12). Alcohol accumulates in this region as it enables a combination of both favourable hydrophobic and polar interactions. Visualisation of the membrane parallel to the z axis provided an easy means by which to assess where solution species were falling on the membrane surface, and how the bilayer was responding to this (Figure 5(c)). Visualisation of the membrane alone (Figure 5(c1)) showed that there was a lateral separation of the head group regions which exposed the hydrophobic tails to the bulk solution. Including alcohol oxygen atoms within 10 Å of the membrane (Figure 5(c2)) showed that they enter the exposed regions, aligning with the tails just below the head groups. In the case of the water oxygen atoms (Figure 5(c3)), these are located over the head group areas, forming headgroup ‘water caps’, as also shown *via* contact analysis (Figure 4(c-e), Supplementary Figure 12(c-e)). This supports the mechanism proposed and shows that during early stages of deformation significant amounts of water did not enter the membrane. Visual inspection showed that water pore formation in both the SaCM and EcIM was fleeting and rare despite the extensive penetration of alcohol and significant deformation of both membranes.

Measurement of the SASA of CHX alone revealed no significant aggregation (see Supplementary Figure 15). CHX was again seen to adopt a c-shape binding (see Supplementary Figure 11). This mode of binding is especially favourable as its polar termini can form interactions with the phosphate/ester region of the lipids at the headgroup-tail interface. Although the ‘wedge’ conformation proposed by Komljenović *et al*[18] was observed on occasion, it was far less common than the c-shape conformation. The results presented here for PL membranes are strongly supported by the findings of Rzycki *et al*.[33] who also observed CHX molecules may bind individually rather than in clusters, and those which did bind as an aggregate would rapidly disperse. Evidence of this dispersal is shown in Supplementary Figure 9. Similarly, they also found CHX to bind preferentially in the c-shape conformation and that the effect of CHX was minimal.

### Electroporation insertion of CHX and effect of gluconate: MD study of the action on the EcIM

To eliminate the possibility that membrane permeation of CHX into the EcIM was not observed due to insufficient simulation sampling, we initiated simulations with CHX already placed in the core of the membrane. An external, constant electric field (0.125 V nm^-1^) was applied across the EcIM to create a large water-filled pore. A CHX molecule was manually placed within this pore and the electric field was then turned off to determine whether CHX would remain in the membrane core as the pore closed (as in the work of Piggot *et al*.[34]) Over the course of a 200 ns production run, CHX shifted from the membrane core to the same location as observed in the equilibrium simulations of CHX presented above (see Supplementary Figure 16). This behaviour was observed across all three of the independent simulations of this system, providing conclusive support that headgroup-lipid interface is the preferred location for CHX.

CHX is usually combined with gluconate (GLUC) in sanitising agents for increased solubility. To eliminate the possibility that GLUC is required in order to observe deep membrane penetration of CHX, we performed simulations including both molecules. 20 GLUC molecules were thus added to the bulk water (0.5% w/v) phase surrounding the EcIM (see Supplementary Figure 17). Following 100 ns of production simulations across three repeats, GLUC was not observed to penetrate the membrane and only briefly interacted with the surface (< 1 ns events). Additional simulations with 10 CHX molecules were added to the same system showed the same CHX membrane binding mode as observed in simulations without GLUC. GLUC molecules were observed to regularly bind to the CPL regions of CHX, but this did not change the binding mode. (see Supplementary Figure 18).

### Chlorhexidine in alcohol solutions: action on the *E. coli* outer membrane (Ra-LPS)

The action of the chlorhexidine/alcohol solutions on the EcOM was nex studied in the same way as for the PL membranes. The EcOM was extremely resistant to the deformation caused by alcohol in simulations of the PL membranes, despite the alcohol distributing within the membrane in the same way as for PL bilayers.[33] This is likely due to the strong interactions between LPS molecules and its stabilising ‘cross-linking’ calcium counter ions. Over the course of the first 60 ns (200 ns simulations in triplicate) there was no discernible change in the membrane structure in any of the EcOM simulations. By this time both PL membranes (SaCM and EcIM) had shown significant deformation due to alcohol partitioning in the headgroup/tail interface. Due to the retained integrity of the EcOM there was little change when analyzing the bilayer density, other than a small shoulder of budding lipids (see Supplementary Figure 19). By 80 ns the PL leaflet of the EcOM began to form a small bundle of approximately 50 lipids which started to separate from the membrane (Figure 6(a)). The budding phenomenon was observed in two of the three repeats with PROH but none of the repeats performed in ISOP (see Supplementary Figure 20). PROH distributes into the lipid headgroup/tail interface of both leaflets, whereas ISOP did not distribute into the interface as effectively because of its branched structure. Partitioning of alcohol into the membrane disperses the bilayer components due to crowding. This was alleviated in the PL membranes by lateral dispersal, however due to the aforementioned calcium cross-linked LPS molecules lateral movement was prevented here. This resulted in the PL leaflet budding into the periplasmic space to relieve the crowding, forming a shell of polar lipid headgroups sheltering their hydrophobic tails. As a small amount of both alcohols was able to pass through the LPS leaflet regularly, this showed that deformation is possible with alcohol entering from only the extra-cellular leaflet, as expected physiologically. Given enough time to penetrate the membrane solely from the LPS leaflet side, build up of lateral pressure may result in budding to such a degree that it entirely deforms the EcOM. As commercial sanitizers contain more alcohol (∼70% w/v) this may be expected to occour at an even faster rate than observed here.

**Figure 6.**
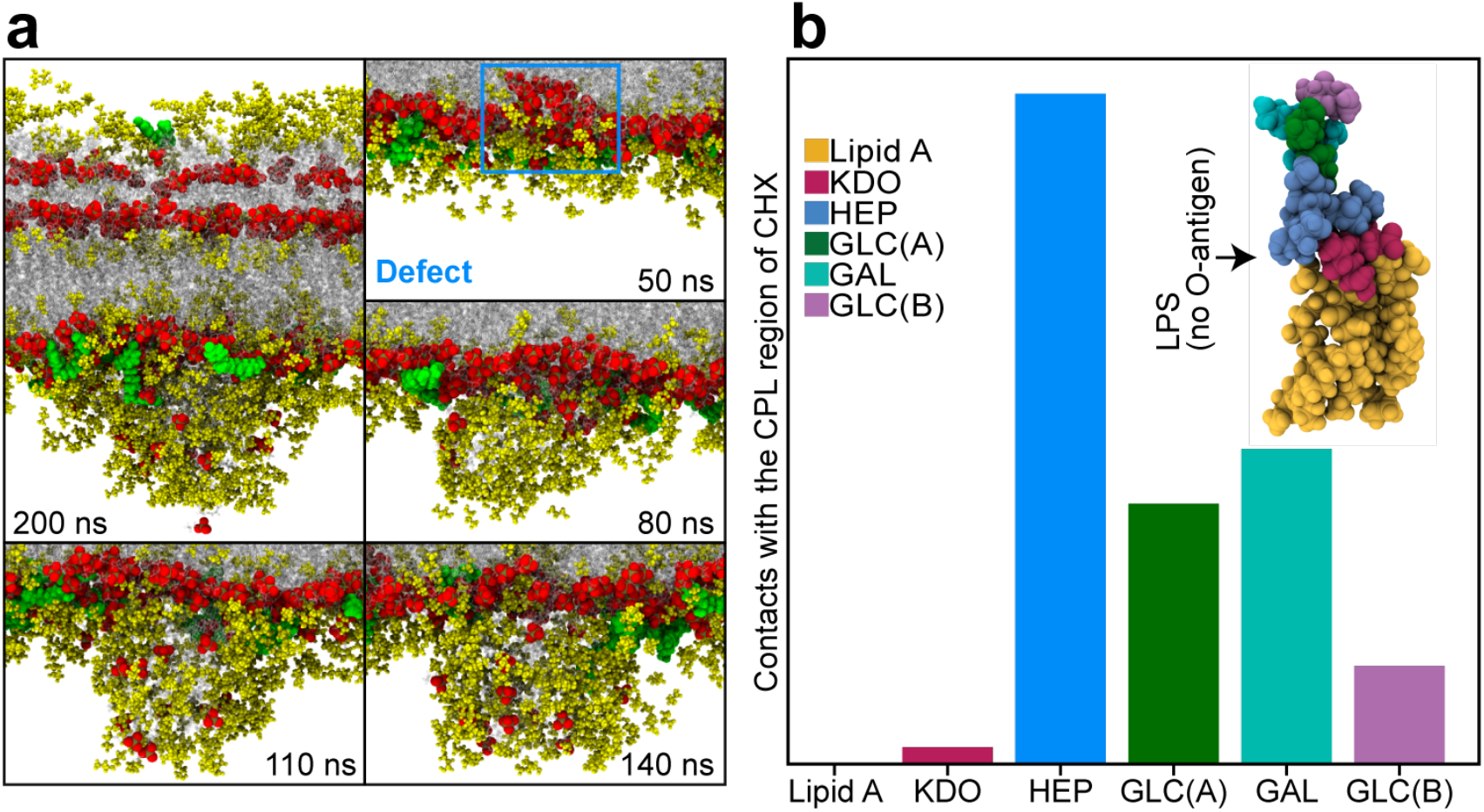
(a) Snapshots of the EcOM exposed to 0.5% w/v CHX in 20% PROH at 200 ns with further snapshots of the PL leaflet at 50, 80, 110 and 140 ns, showing phosphate oxygen as spheres in red, phosphate phosphorous in gold, CHX in green, PROH within 10 Å of the membrane in yellow and lipids as translucent. (b) The proportion of contacts between the CPL region of CHX and sugars in LPS tracked over the full 200 ns of production by checking at each time step how many contacts there were within 2.5 Å of the CPL region and adding them to the total count per sugar, showing lipid A in yellow, ketodeoxyoctonic acid (Kdo) in red, Heptose (HEP) in blue, Glucose (GLC) A in green, Galactose (GAL) in cyan and Glucose B in pink. Glucose are labelled A and B as they are in two different positions.

Visual analysis of the production runs showed that only one CHX molecule ever became bound to the LPS leaflet in any of our simulations and in three of out six simulations with alcohol and CHX present, none became bound to the LPS leaflet (2/3 PROH repeats and 1/3 ISOPR repeat). The lack of binding to the LPS leaflet was due to the significantly fewer favourable interactions between the HEX region of CHX and LPS sugars. Generally CHX became bound to the heptose moeity in the LPS sugar region and did not translocate any further (Figure 6(b)). A summary of the interaction of the LPS leaflet with CHX is shown in Supplementary Figure 21 and Figure 22.

### Chlorhexidine and propanol: ^31^P NMR study of their action upon the structure of the *E. coli* inner membrane

The action of sanitizer components on the structure of the EcIM model was next assessed experimentally using ^31^P solid-state NMR. This method produces a lineshape characteristic of the dynamics and phase of a lipid bilayer. Spectra were acquired of multilamellar vesicles composed of PE/PG/CL (90/5/5 mol%) exposed to PROH, GLUC and CHG to assess their influence on the bilayer structure (Figure 7).

**Figure 7.**
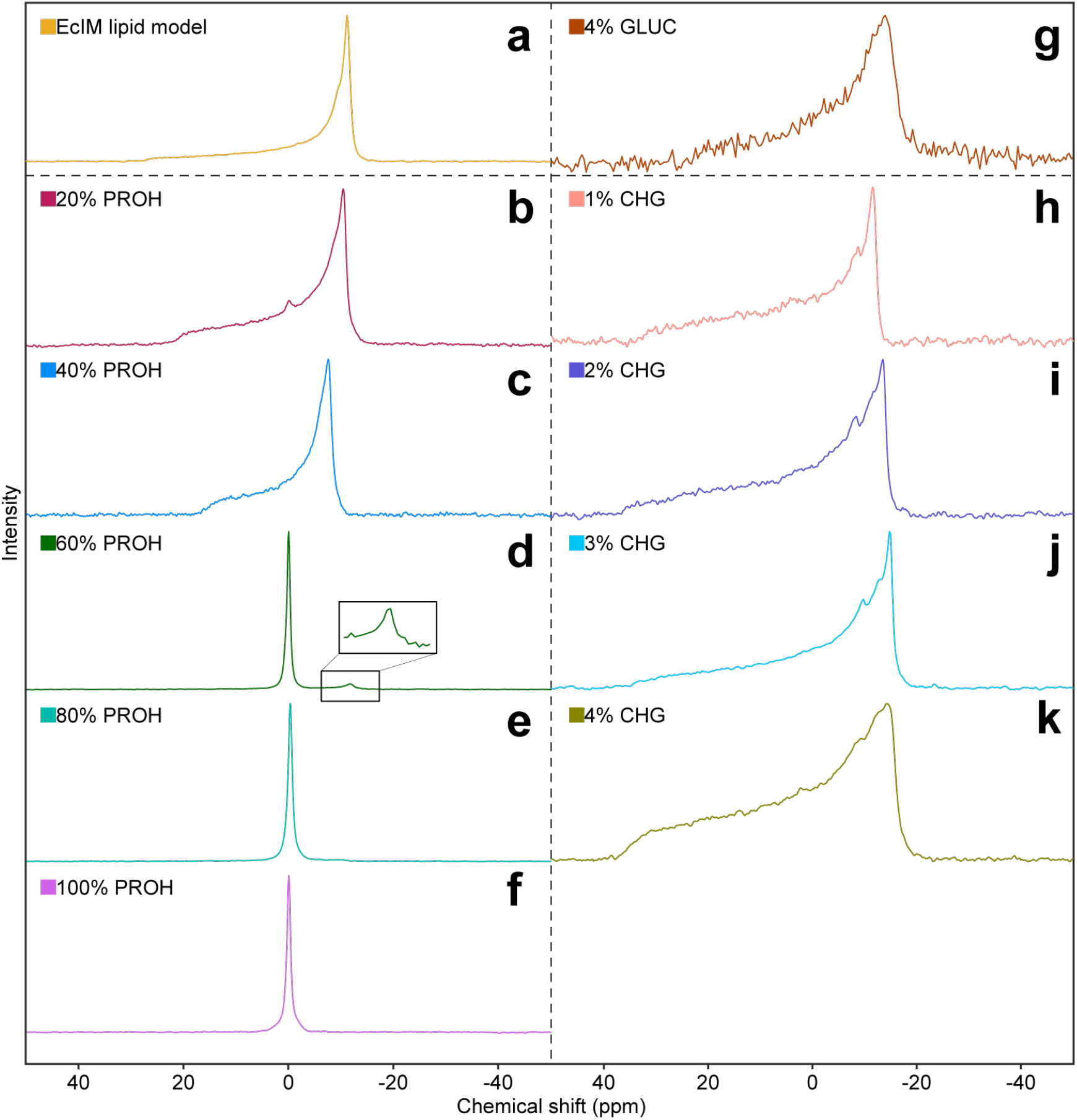
Static ^31^P NMR spectra of the EcIM model (A) alone, with PROH at (B-F) 20% w/v intervals, with (G) 4% w/v GLUC and with CHX (H-K) up to 4% at 1% w/v intervals. The area marked by an asterisk is the emerging environment which was witnessed in the CHX experiments.

As expected in the absence of any reagents, the ^31^P spectrum of the model EcIM exhibited an axially symmetrical lineshape, with a chemical shielding anisotropy of 43 ppm, consistent with the lipids existing in the liquid crystalline phase. Increasing the concentration of PROH in the sample resulted in significant perturbations of the bilayer structure (Figure 7(b-f)). Up to 40% the lipids retained their bilayer character, with a slight reduction in the chemical shielding anisotropy indicative of increased headgroup mobility. This was consistent with the partitioning of the PROH at the top of the lipid chains that would result in a lower lateral pressure in the headgroup region. Between 40% and 60% w/v PROH most of the bilayer structures were solubilised with the spectra dominated by an isotropic signal at -0.1 ppm, with only a minor bilayer-like component still present at 60% PROH.

To assess the influence of CHG on the bilayer structure the ^31^P lineshape was measured at concentrations up to 4% w/v (Figure 7(h-k)) as this is the highest concentration commonly used in commercial sanitizers. Increasing concentrations of CHG had limited influence on the ^31^P lineshape, with all spectra showing a characteristic axially symmetric lineshape indicative of a bilayer like phase. Notably there was a slight increase in the chemical shielding anisotropy as the concentration increased, suggesting that the CHG may limit headgroup mobility. This was also reflected in subtle changes in the lineshape. Increases in the intensity on the downfield edge and overall broadening of the spectral features were similar to those observed in the gel phase and which are typically attributed to a reduction in T2 of the phosphates in the headgroup. This suggests that the bilayer became more rigid and less elastic, likely due to CHG binding holding lipid headgroups together, as observed *via* MD simulation.[35–39]

Similar observations were seen upon the addition of GLUC with the spectra (Figure 7(g)) exhibiting similar broadening due to a reduction in T2, although no changes in the relative intensity across the lineshape were observed, thereby suggesting that GLUC had minimal influence on bilayer elasticity. These effects appeared to be mediated by CHX, as the corresponding spectrum of 4% GLUC exhibits a classical axially symmetric lineshape expected from multi lamellar vesicles (MLV).

### Chlorhexidine and propanol: an ^1^H MAS-NMR study of their location within the *E. coli* inner membrane

To investigate the location of the CHG within the model EcIM, ^1^H-MAS and 2D ^1^H/^1^H MAS-NOESY experiments were recorded. Due to the high mobility within the lipid bilayer the ^1^H spectra were well resolved, as previously reported.[14,40] The ^1^H spectrum of the model EcIM was dominated by resonances from POPE, the major lipid species in this system. The resonances from the lipids were assigned based on published assignments of POPC and POPE (Figure 8).[14] To assist in the assignment of the components present in sanitizer, the individual PROH, GLUC and CHG components were added to the EcIM independently. Addition of 20% PROH gave rise to three major resonances at 3.61, 1.61 and 0.96 ppm whilst the addition of GLUC gave rise to a family of resonances superimposed on lipid signals assigned to G1, G3, α and β. Assignments were made based on published values and are shown in Figure 8^30^. The addition of CHG resulted in the appearance of a further 6 resonances which were assigned to CHX, with those in the aromatic structure clearly resolved (Figure 8(d) - proton assignments 1 and 2), whilst those from the BGU/HEX appearing amongst the resonances arose from the lipids in the sample (Figure 8(d) - proton assignments 3,4 and 5). Closer inspection showed that resonances arising from the C2, C3 and G2 sites within the lipid were shifted upon the addition of CHG. It is plausible that the presence of the CPL groups at the headgroup-tail boundary may have generated ring-current effects resulting in the observed perturbations, in a manner similar to that reported for ibuprofen by Kremkow *et al*.[41] Close inspection of the CPL region of the spectrum (Figure 8(f)) revealed that CHG exhibited three resonances in the aromatic region, which was unexpected given the presence of only two resonances for the aromatic protons in the CPL groups in CHX and the expected magnetic equivalence of the protons on either side of the aromatic ring. We suggest that the additional resonance arose due to a stable complex of CHX and GLUC, formed by interaction of the CPL region of CHX with the carboxylic acid group of GLUC (Figure 8(g)). This would break the magnetic equivalence and perturb the C2 shift in a manner similar to that observed. This was supported by the complexes seen in MD simulations of CHX with GLUC (Figure 8(e)).

**Figure 8.**
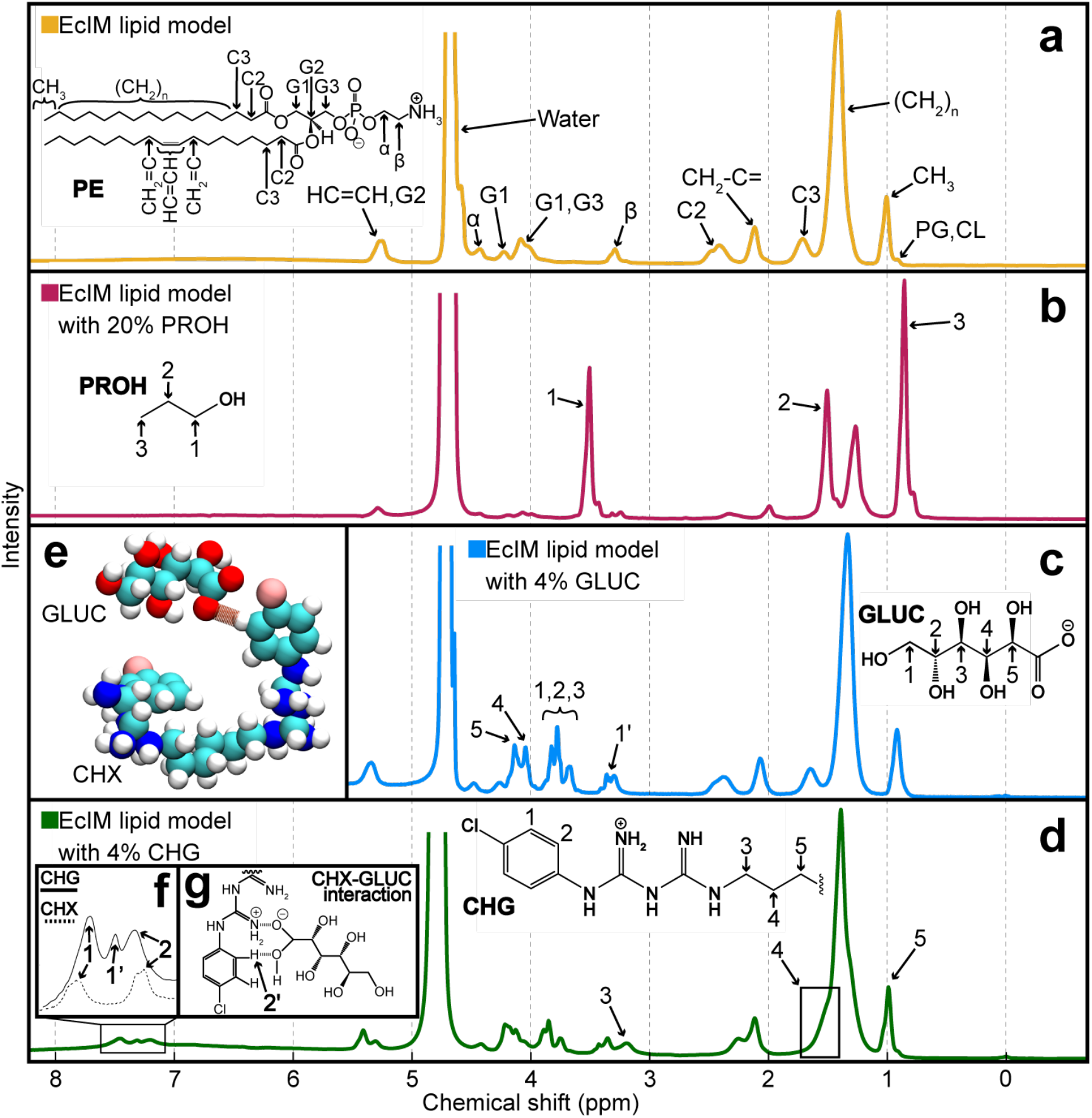
Assignment of the ^1^H NMR spectra of (A) the EcIM model alone in water, (B) with 20% w/v PROH, (C) with 4% w/v GLUC and (D) with 4% w/v CHG. (E) A visual representation of a common interaction between CHX and GLUC in an MD simulation of 10 CHX and 20 neutralising GLUC. Atoms are shown as spheres, hydrogen in white, carbon in cyan, nitrogen in blue, chlorine in pink, oxygen in red and hydrogen bonding in brown. (F) Resonances arising from CHX and CHG. (G) A mechanistic depiction of the interaction between CHX and GLUC which is theorised to account for the additional resonance.

### Chlorhexidine: a ^1^H NOESY MAS-NMR study of their location within the *E. coli* inner membrane

To establish a more detailed idea of the average location of the CHX within the membrane, 2D ^1^H/^1^H NOESY MAS-NMR spectra were recorded. The dynamics present in the membrane give rise to the well resolved ^1^H spectra, at moderate MAS frequencies, allowing the analysis of NOE based magnetization exchange. The intensity of the cross-peaks in the 2D ^1^H/^1^H NOESY MAS-NMR reflect the residency time and proximity of protons with respect to one another.[42] As shown in previous research,[42–44] an analysis of the intermolecular cross-relaxation rates between resonances arising from the small molecule and the lipids can provide a valuable insight into the partitioning and location of small molecules within the lipid bilayer. CHG was applied to the membrane at a concentration of 4% w/v (Figure 9). Although the percentage volume was much larger, due to the size of CHG the molar ratios of PROH and CHG to lipids are similar (1:4.67 and 4.12 respectively). There were significant intramolecular cross-peaks between the resonances arising from the CHG, most clearly defined with sites in the CPL rings and less clearly between the HEX resonances which were poorly resolved from lipid resonances. Inter-molecular cross-peaks were also apparent; these were most clearly observed between the CPL protons and the intense signal from the CH2 moieties in the lipid chains. Lower intensity peaks were also apparent between the CPL protons and those from the protons at C2 and the glycerol backbone. This data again indicates that CHG has a preferred localisation at the level of the glycerol backbone in the lipid bilayer. Weak correlations were also seen between the C3 and the lipid resonances, indicating that the backbone of the CHG was lying in the plane of the membrane, in agreement with the MD simulations presented above. To ascertain the relative orientation of CHG within the EcIM, a quantitative analysis of the cross-relaxation rates was performed (Figure 8(b)). Interestingly, the cross-relaxation rates suggest that resonances 1 and 2 interacted favourably with sites spanning the region from the unsaturated site within the lipid chains (HC=CH), up to the region of the lipid backbone. In contrast, cross-relaxation of resonance 3 was dominated by exchange to the α proton of the PE headgroup, suggesting that the BGU/HEX groups of CHG prefer to localise within the region of the lipid headgroups, in keeping with the membrane bound position observed during simulation.

**Figure 9.**
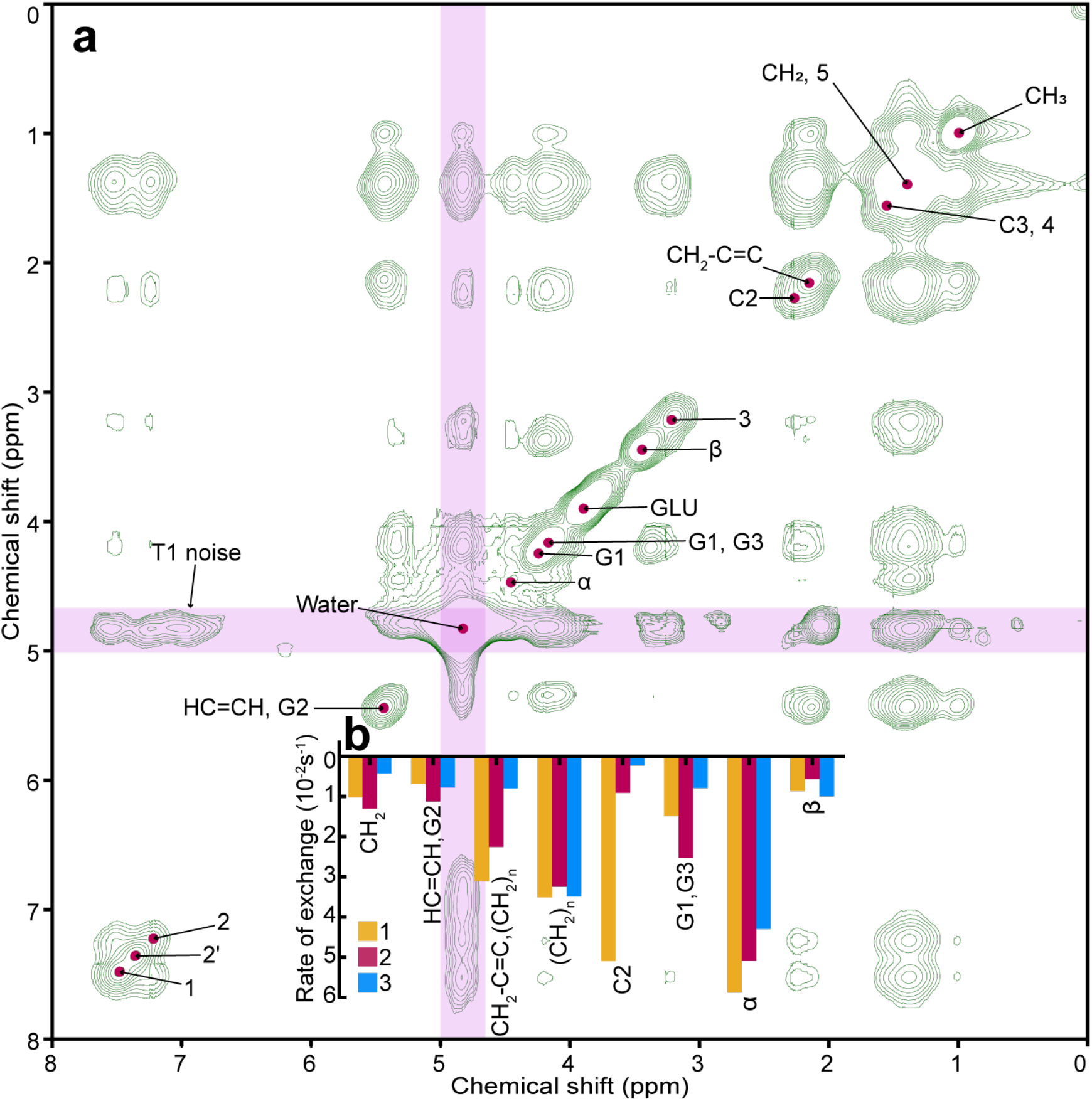
Assignment of 2D NOESY spectrum of the EcIM model membrane when exposed to CHG for calculation of magnetisation exchange rates between protons in the system. (a) The two-dimensional NOESY NMR spectrum of the EcIM model exposed to CHG at a concentration of 4% w/v with a 250 ms mixing time. Positive contours are shown in green and peaks are indicated by a mark in red. Areas which are distorted due to T1 noise are highlighted in pink. (b) A bar chart of the magnetisation exchange rate of the protons 1, 2 and 3 with protons in lipids in the EcIM lipid bilayer model, ordered from left to right to represent groups ascending from the bilayer centre toward the top of the lipid head group.

## CONCLUSIONS

The simulation and NMR studies described here provide insights into the mode of action of the common sanitizing agent CHX on *S. aureus* and *E. coli* membranes. We find that in the aqueous solution (no alcohol), CHX binds and inserts into the PL membranes (SaCM and EcIM) and maintains a position at the headgroup/tail interface of the latter, while moving slightly deeper into the former. Intriguingly, insertion of CHX does not induce disruption of the membranes during the timescales of the MD simulations. Nevertheless, CHX was found to reduce the MSD of lipids due to binding and holding together together headgroups, thereby acting as a ‘molecular staple’. This result also supports a previously suggested mechanism.

No insertion of CHX into the LPS leaflet of the EcOM was observed in the MD simulations. This is likely due to limitations in timescale exacerbated by the slow-moving nature of LPS. We note that similar lack of penetration of other cationic antimicrobial agents such as polymyxins has previously been reported from atomistic MD simulations.[45] Nevertheless, the comparative simulations show a greater kinetic barrier to entry into the outer membrane compared to the PL membranes.

Inclusion of alcohol, PROH or ISOP, has a dramatic impact on the PL membranes. Both alcohols were seen in multiple simulation replicas to cause significant deformation of both the EcIM and SaCM. Although alcohols deformation of PL membranes is not a novel finding, contact tracking has shown that deformation is the product of many transient residencies of alcohol in energetically beneficial positions within the bilayer. Large quantities of alcohol partitioning into this region resulted in lateral dispersion of lipids which caused significant deformation; at larger alcohol concentrations this leads to dissolution of the membrane. The same effect as was seen for PL membranes was not seen for the EcOM due to the lateral interactions between components of the LPS leaflet and calcium which are significantly stronger compared to either PL membranes. Restrictions imposed by the LPS leaflet meant that the bilayer could not disperse to alleviate steric crowding. This caused the PL leaflet to ‘bud’ into the solution as this was the only way to reduce the crowding created by alcohol partitioning.

Solid-state 1D proton NMR studies corroborate the findings of the simulations with specific peaks in the 1D NMR spectra which confirm the CPL groups in CHX bind to the membranes, as predicted computationally. This was then examined in greater detail by exploring the magnetisation exchange rates which were seen for specific protons in CHX. This showed that the binding of CPL in CHX was not only in the same position as observed in simulation, but also adopted the same orientation.

Static ^31^P solid state NMR showed *via* perturbations in lineshape of the MLV that increases in the concentration of alcohol deformed and then dissolved the MLV, as expected. Increasing the concentration of CHX from 1-4% w/v did not lead to destruction of the MLV bilayer. Very minor changes in the line shape, contrarily, suggested an increase in the rigidity of the MLV bilayer. This finding was corroborated by the lower MSD of lipids in the simulated EcIM and SaCM systems when CHX was applied; decreased mobility implies greater membrane rigidity.

These data have successfully begun to elucidate key details regarding the modes of action of sanitizer components when applied to common bacterial membranes. By studying the differences between these systems, we can begin to further develop chemicals to improve their membrane destructive properties, particularly in light of growing antimicrobial resistance. In future studies we aim to include other membrane bound molecules such as proteins, against which sanitizers may also act. A course-grained approach with CHX would also allow us to explore time limitations and what changes may be induced when observed over significantly longer timeframes. Broadening our approach to these systems will allow us to further assess what is currently limiting the sanitizers we currently rely upon and how bacteria may evolve to counter them.

## METHODOLOGY

### Simulation system setup and model parameters

Aqueous alcohol solutions were constructed in the desired percentage (20%) before performing equilibration and production. Membranes were surrounded by an aqueous KCl (0.15 M) solution before equilibration and production. Alcohol solutions were applied to both faces of the membrane by removing water from the membrane system and adding the aqueous alcohol solution in the cleared space, without removing KCl from the system.

### Simulation protocols

Simulations were set up and performed using the GROMACS[46–48] molecular dynamics software package (version 2020.2) with the CHARMM36 force field.[49,50] Systems were maintained at a temperature of 310 K, using the Nosé-Hoover[51,52] thermostat with a time constant of 1 ps. The pressure of the system was maintained at 1 atm, with a time constant of 2 ps, using semi-isotropic pressure coupling with the Parrinello-Rahman[53,54] barostat. All van der Waals interactions were cut off at 1.2 nm and a smooth particle mesh Ewald[55] (PME) algorithm was used to treat electrostatic interactions with a real space cut off of 1.4 nm. Simulation parameters were chosen based on similar published studies of Piggot et al.[34] For equilibration, each system was subjected to 500 ps of NVT simulation, followed by 2 ns of NPT. Positional restraints (1000 kJ mol^-1^ nm^2^) were placed on the membrane head group atoms during NVT and NPT equilibration. Production simulations were then performed without any positional restraints. The results were analysed using GROMACS tools and in-house scripts. Visualisation was performed using the visual molecular dynamics (VMD)[56] software package. Electroporation of the membrane was performed with a 0.125 V nm^-1^ electric field applied along the membrane normal.

### Calculating proportion of sugar interactions with the CPL region of CHX

The full 200 ns of production was parsed through a python script written in Jupyter Notebooks[57] which checked each timeframe for contacts within 2.5 Å of the CPL region *via* MDAnalysis[58] and then used this information to create a bar chart with Matplotlib.[59]

### Experimental Sample Preparation

Lipid samples were prepared from a stock solution of PE, PG and DPG in a ratio of 90:5:5 respectively. Stocks were placed in ethanol at a lipid concentration 10 mg ml^-1^. Samples of 5mg lipid mixtures were left under high-vacuum to remove residual solvent. The lipids were rehydrated in 20 µL of D2O (25% w/v) containing alcohol or chlorhexidine at the appropriate concentration. Samples containing 5 mg of lipid were freeze-thawed 5 times. Each cycle involved placing the sample in liquid nitrogen, thawing and mixing with a vortex mixer. These samples were then transferred to 3.2 mm rotors.

### 600 MHz solid-state NMR

Solid-state NMR spectra were recorded on a 600 MHz Agilent DD2 NMR spectrometer (Yarnton, UK), equipped with a 3.2mm triple resonance MAS NMR probe. All spectra were acquired at 25 °C unless specified. Static ^31^P spectra were recorded with a Hahn-echo pulse sequence[60] with a 3.5 ms excitation pulse, 70 kHz proton decoupling during acquisition and a 50 ms echo time. A 2.5 second recycle delay was employed to minimise sample heating. All ^31^P spectra were externally referenced to H3PO4 (85%). Prior to Fourier transformation data was left shifted to the top of the echo, zero filled to 2048 points and 100 Hz linebroadening was applied.

All proton spectra were record with 12.5 kHz MAS, and a 2.75 ms pulse for excitation, and a 2.5 second recycle delay. All proton spectra were acquired with the probe with the X channel tuned to deuterium and connected to the spectrometers lock allowing the stabilization of the field which significantly reduced t1 noise in 2D spectra. Proton spectra were referenced to the residual water peak at 4.65 ppm. 2D ^1^H-^1^H MAS-NOESY spectra were recorded using a standard exchange sequence[61] with States-TPPI in the indirect dimension.[62] Data was processed with in NMRpipe[63] or Matlab[64] using matNMR.[65]

### Matrix based approach to NOESY magnetisation exchange rates

Using the two-dimensional NOESY spectra acquired, the rate of exchange of magnetisation for protons in the system were calculated *via* a full matrix approach based on the Solomon equations.[43] Peak volumes were integrated using the ccpNMR software.[66] The resulting volume matrices were then normalised and used to calculate the cross-relaxation rates according Supplementary Equation 1 using custom written scripts in Matlab (Mathworks).

## Abbreviations

CHX: Chlorhexidine
CPL: Chlorophenyl
BGU: Biguanide
HEX: Hexane
APL: Area Per Lipid
SaCM: *S. aureus* cell membrane
EcIM: *E. coli* inner membrane
EcOM: *E. coli* outer membrane
LPS: Lipopolysaccharide
PE: Phosphoethanolamine
PG: Phosphoglycerol
DPG: Cardiolipin
LPG: Lysyl-Phosphatidylglycerol
SASA: Solvent Accessible Aurface Area
PROH: Propanol
ISOP: Isopropanol
GLUC: Gluconate
VMD: Visual Molecular Dynamics
NOESY: Nuclear Overhauser Effect Spectroscopy

