## Supplementary for "Impact on *S. aureus* and *E. coli* membranes of treatment with chlorhexidine and alcohol solutions: insights from molecular simulations and nuclear magnetic resonance"

Declarations of interest: None

### Supplementary Information

#### Chlorhexidine in water

CHX is known to have poor miscibility in water so is usually utilised as CHG in sanitizers. 10 CHX molecules were simulated at two different concentrations of KCl (0.15 and 1 M) for 200 ns. Supplementary Figure 23(a) shows that the larger concentration of ions results in molecules having a much smaller average radial distribution function (RDF). The decrease in distance is caused by an increase in interactions between CHX molecules mediated by ions. Supplementary Figure 23(c-e) provide visual analysis of key time steps when SASA (see Supplementary Figure 23(b)) was low. CHX molecules in the system came together to form large clusters which coincided with significant drops in SASA. SASA studies were performed on extracted trajectories of only CHX, so all changes are due to conformational changes or clustering. With a higher concentration of ions the charged and polar regions of CHX have a larger availability of nucleation points which they can begin to aggregate around, resulting in larger clusters. Dips in SASA show that these clusters do not persist for more than 15 ns. The frequency of occurrence of the most common clusters was reported to determine if the clusters were transient or stable complexes (see Supplementary Figure 24). CHX in 0.15 M salt was more likely to result in molecules in one of the 5 most common clusters, while 1 M salt systems molecules were distributed over a larger number of clusters. Due to the larger concentration it became more likely for CHX to take a wider variety of conformations. To expand upon this the 5 most common conformations of CHX in both systems were extracted and reported with the percentage of conformations they accounted for over the 200 ns production (see Supplementary Table 3). Conformations differ mainly by the orientation of their BGU-CPL region from the HEX functional group, suggesting that these rotate to accommodate more energetically favourable interactions. This is consistent with the theoretical binding mechanism reported for CHX. Simulations of CHX were performed at 0.15 M as this is closer to physiological concentration and will better reflect phenomenon which may be expected *in-vivo*. Due to an already constricted time frame that these simulations can cover it makes no sense to use a higher concentration if it is likely that clustering will lessen the availability of CHX to act on the membrane.

| Mem. | Alc. | | CHX | | GLUC | | KCl conc. / M | E. field / V nm^-1^ | Reps. / ns |
| --- | --- | --- | --- | --- | --- | --- | --- | --- | --- |
|  | **Type** | **Conc. / vol. %** | **No.** | **Conc. / vol. %** | **No.** | **Conc. / vol. %** |  |  |  |
| None | - | - | 10 | 0.5 | - | - | 0.15 | - | 3 x 200 |
| None | - | - | 10 | 0.5 | - | - | 1.00 | - | 3 x 200 |
| SaCM | - | - | 10 | 0.5 | - | - | 0.15 | - | 3 x 200 |
| EcIM | - | - | 10 | 0.5 | - | - | 0.15 | - | 3 x 200 |
| EcOM | - | - | 10 | 0.5 | - | - | 0.15 | - | 3 x 200 |
| SaCM | PRO | 20 | - | - | - | - | 0.15 | - | 3 x 200 |
| SaCM | ISOP | 20 | - | - | - | - | 0.15 | - | 3 x 200 |
| EcIM | PRO | 20 | - | - | - | - | 0.15 | - | 3 x 200 |
| EcIM | ISOP | 20 | - | - | - | - | 0.15 | - | 3 x 200 |
| EcOM | PRO | 20 | - | - | - | - | 0.15 | - | 3 x 200 |
| EcOM | ISOP | 20 | - | - | - | - | 0.15 | - | 3 x 200 |
| SaCM | PRO | 20 | 10 | 0.5 | - | - | 0.15 | - | 3 x 200 |
| SaCM | ISOP | 20 | 10 | 0.5 | - | - | 0.15 | - | 3 x 200 |
| EcIM | PRO | 20 | 10 | 0.5 | - | - | 0.15 | - | 3 x 200 |
| EcIM | ISOP | 20 | 10 | 0.5 | - | - | 0.15 | - | 3 x 200 |
| EcOM | PRO | 20 | 10 | 0.5 | - | - | 0.15 | - | 3 x 200 |
| EcOM | ISOP | 20 | 10 | 0.5 | - | - | 0.15 | - | 3 x 200 |
| EcIM | - | - | - | - | 20 | 0.5 | 0.15 | - | 3 x 200 |
| EcIM | - | - | 10 | 0.5 | 20 | 0.5 | 0.15 | - | 3 x 200 |
| EcIM | - | - | 10 | 0.5 | - | - | 0.15 | 0.125 | 3 x 200 |

**Table 1.** The words membrane (mem.), alcohol (alc.), concentration (conc.), volume (vol.), number of molecules (no.), electric field (E. field), repeats (reps.) and section (sec.) are abbreviated in the headings.

| Sanitizer component | Sanitizer : lipid  X : 10 | Concentration / % (w/v) | Experiments performed |
| --- | --- | --- | --- |
| - | N/A | N/A | 1D ^1^H, 1D ^31^P, 2D ^1^H NOESY |
| CHX | 1.00 | 2.46 | 1D ^1^H, 1D ^31^P, 2D ^1^H NOESY |
| CHG | 0.57 | 1 | 1D ^1^H, 1D ^31^P |
| CHG | 1.14 | 2 | 1D ^1^H, 1D ^31^P |
| CHG | 1.70 | 3 | 1D ^1^H, 1D ^31^P |
| CHG | 2.27 | 4 | 1D ^1^H, 1D ^31^P, 2D ^1^H NOESY |
| PROH | 7.70 | 20 | 1D ^1^H, 1D ^31^P |

**Table 2.** To assist the reader, a table of all NMR experiments performed on the EcIM model bilayer is provided above.

| Conc. / M | 1^st^ | 2^nd^ | 3^rd^ | 4^th^ | 5^th^ |
| --- | --- | --- | --- | --- | --- |
| 0.15 | 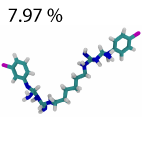 | 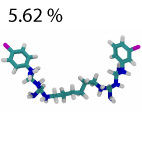 | 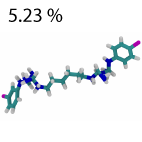 | 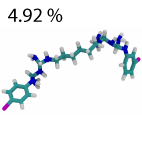 | 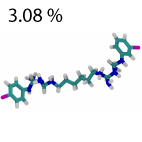 |
| 1.00 | 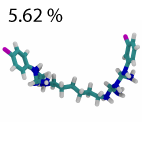 | 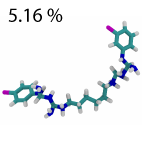 | 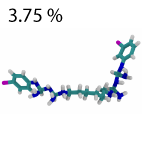 | 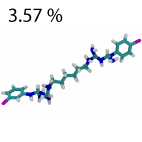 | 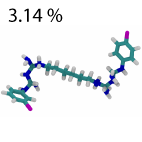 |

**Table 3.** Representation of the five most common conformations of CHX which were seen in cluster analysis of the 0.15 and 1 M KCl systems (see Supplementary Figure 1). Showing carbon in cyan, nitrogen in blue, hydrogen in white and chlorine in purple. The word concentration (conc.) has been abbreviated in the heading.

| Peak | T_1_ relaxation time / s | |
| --- | --- | --- |
|  | ***E. coli*** | ***E. coli* + CHX** |
| HC=CH/G2 | 0.7559 | 0.8989 |
| Water | 3.7600 | 4.3460 |
| α | 1.2340 | 0.6998 |
| G1 | 0.9685 | 0.6627 |
| G1/G3 | 0.8009 | 0.7041 |
| β | 1.0250 | 0.6671 |
| C2 | 0.7007 | 0.6431 |
| CH_2_-C=C | 0.7386 | 0.6591 |
| C3 | 0.6616 | 0.6457 |
| CH_2_ | 0.7820 | 0.6903 |
| CH_3_ | 1.0690 | 0.8704 |
| CHX(1) | - | 0.8575 |
| CHX(2) | - | 0.8353 |
| CHX(3) | - | 0.8108 |
| * | - | 1.7820 |

**Table 4.** T_1_ relaxation times of individual peaks in the EcIM lipid model membrane systems where it is alone, with CHX, with PROH and with both at the same time.

**Figure 1.** The SASA of all CHX molecules combined in the (yellow) SaCM, (red) EcIM and (blue) EcOM systems exposed to CHX in 0.15 KCl solution with no alcohol. Visual snapshots depict transient CHX interactions at the 37 ns for the SaCM, 112 ns for the EcIM and 87ns for the EcOM to show surface interactions. Showing PL head group phosphates with phosphorous in gold, oxygen in red and CHX coloured by residue number to distinguish separate molecules.

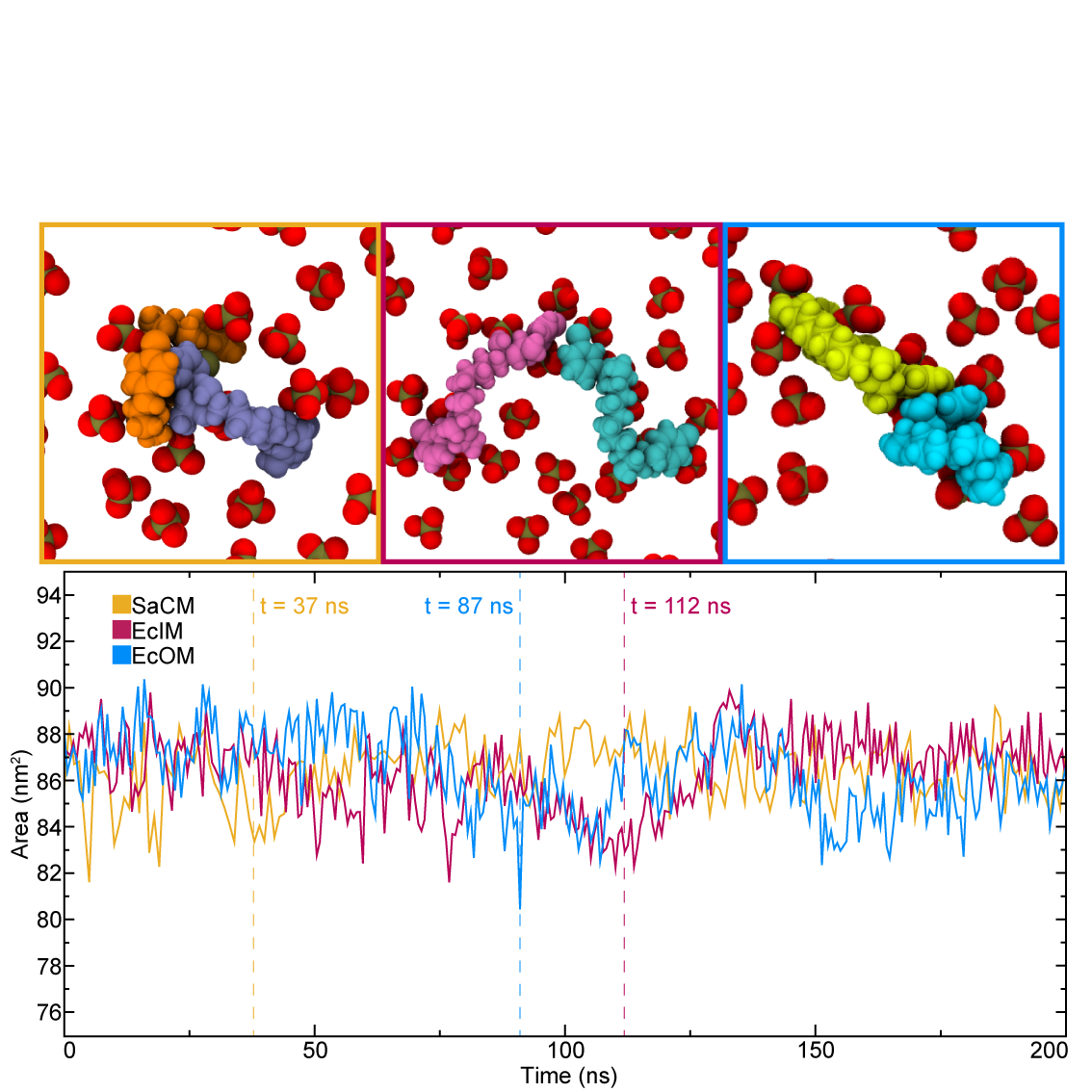

**Figure 2.** The density of membrane components along the z-axis of the SaCM (top), EcIM (middle) and EcOM (bottom) systems exposed to CHX in 0.15 M KCl solution, at (left) 0 and (right) 200. Densities are averaged over 20 ns.

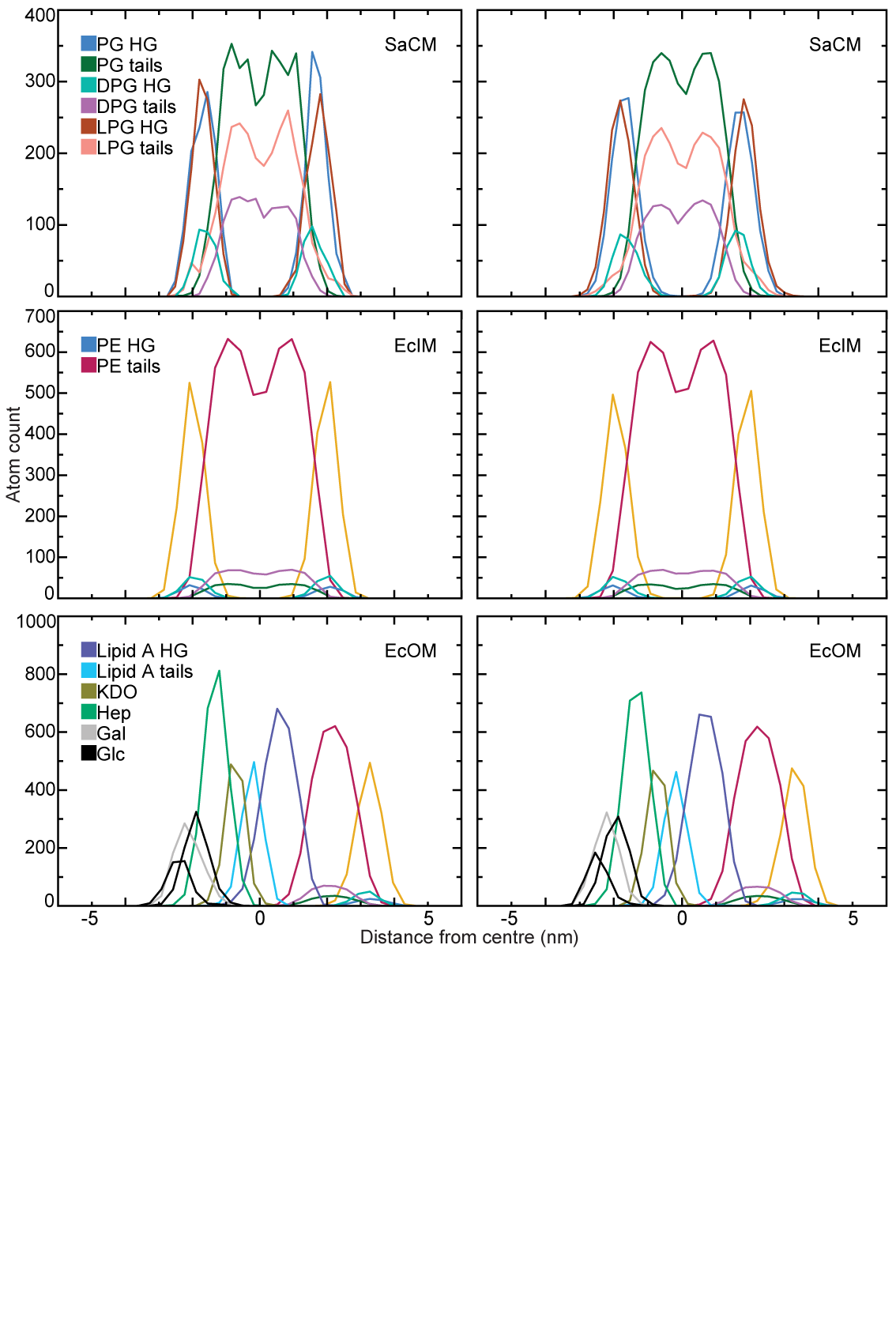

**Figure 3.** Comparison of the average PL APL over 100 ns production runs for the SaCM and EcIM and EcOM systems in aqueous PROH with and without CHX in 0.15 M KCl.

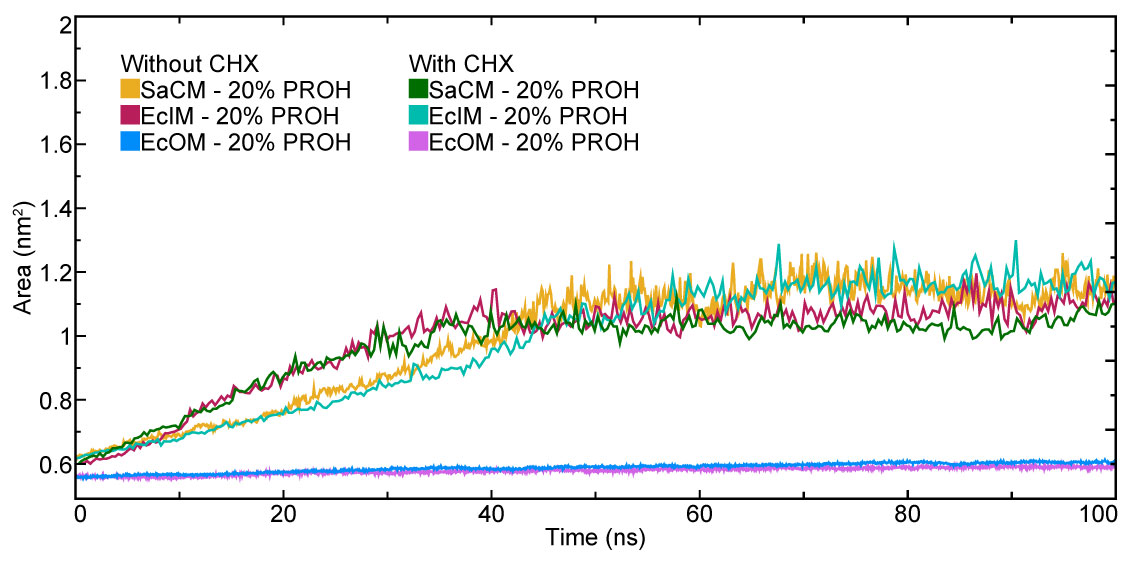

**Figure 4.** Comparison of the average PL APL over 100 ns production runs for all repeats of the EcIM systems in aqueous PROH (yellow) with and (red) without CHX.

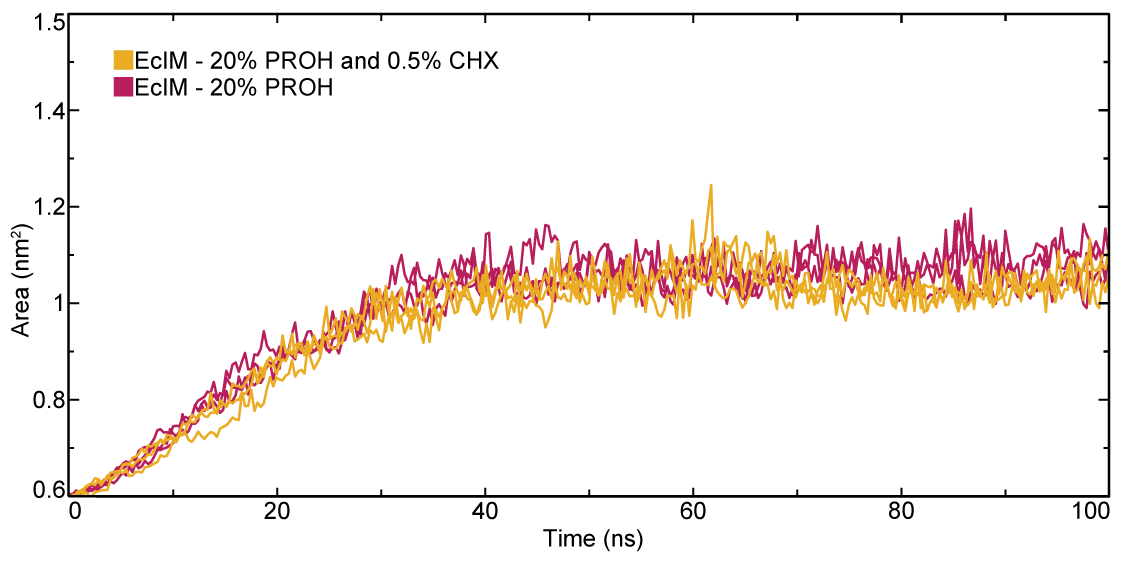

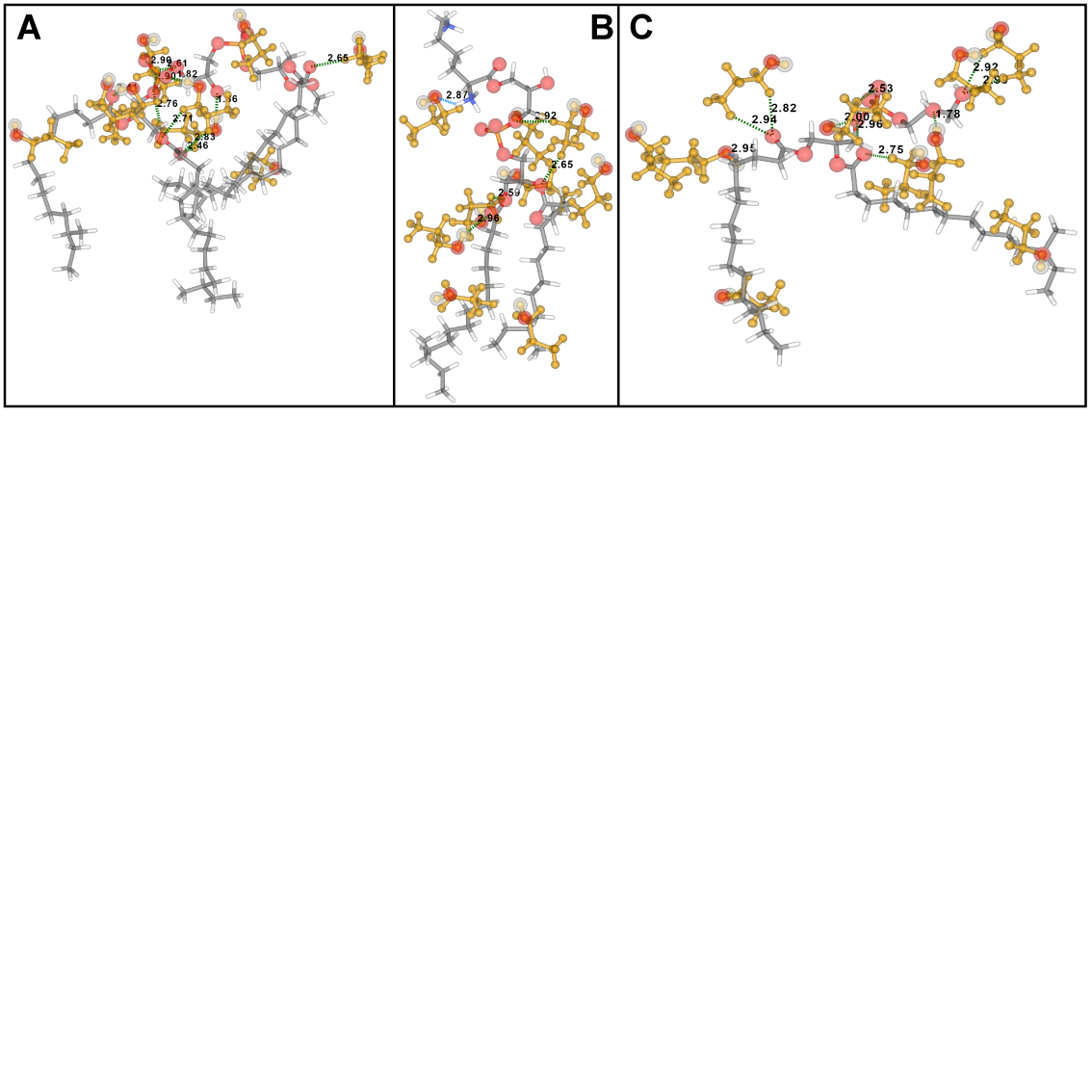

**Figure 5.** A representation of hydrogen bonding interactions under 3 Å between PROH and (A) PG, (B) LPG and (C) DPG in the SaCM, showing hydrogen bonds where PROH acts as the donor in green and acceptor in blue.

**Figure 6.** Density analysis along the z-axis of the SaCM system exposed to (A) PROH and (B) ISOP solutions. Analysed at time intervals (1) 0-20 and (2) 80-100 ns during a 200ns production run.

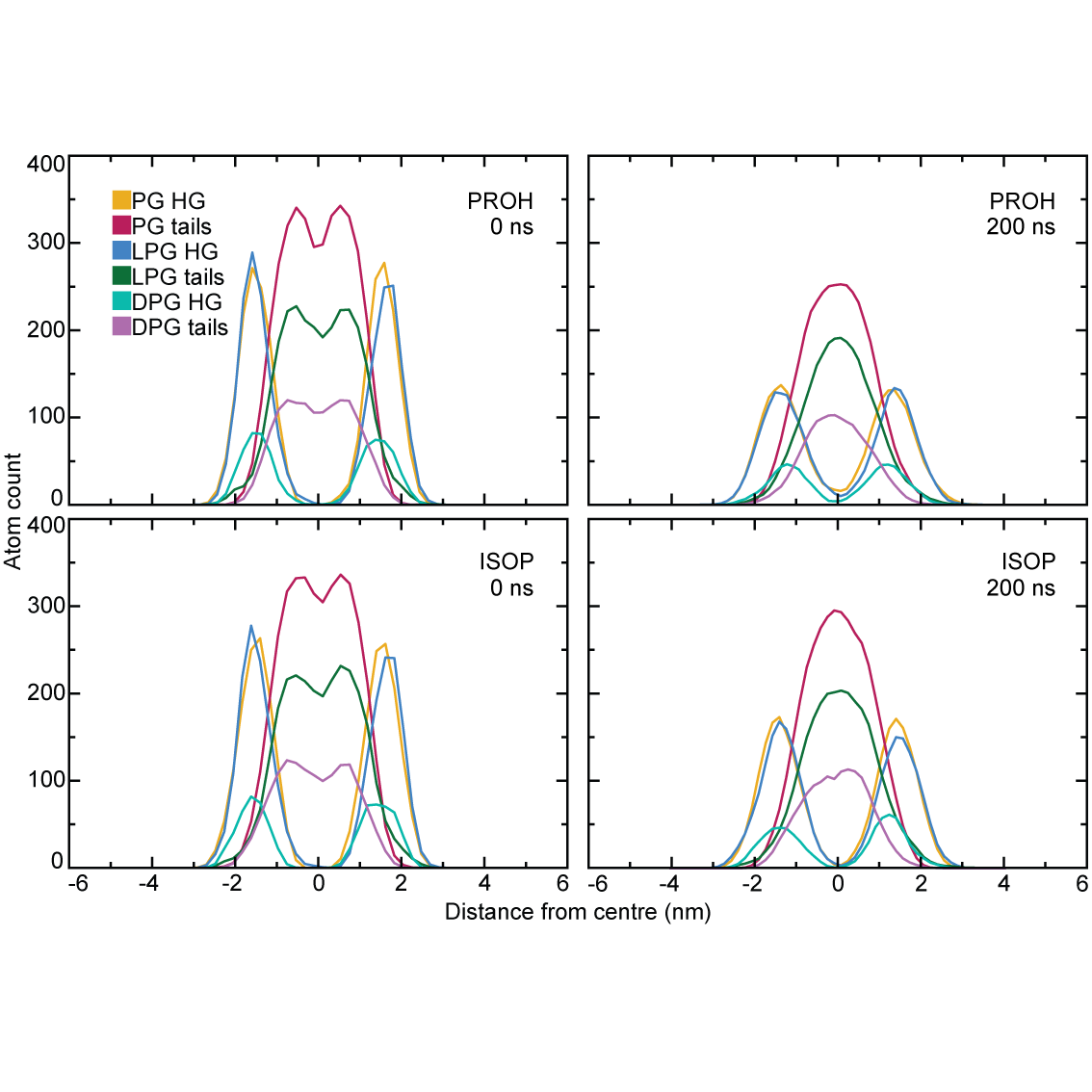

**Figure 7.** The minimum distance between 5 PROH molecules and the (yellow) headgroup, (red) sub-headgroup and (tail) regions of PG in the SaCM over the course of a 200 ns production run. Showing the lower bound hydrogen bond distance in green and periods where the molecule remains within the range of 6 Å of the lipid in cyan.

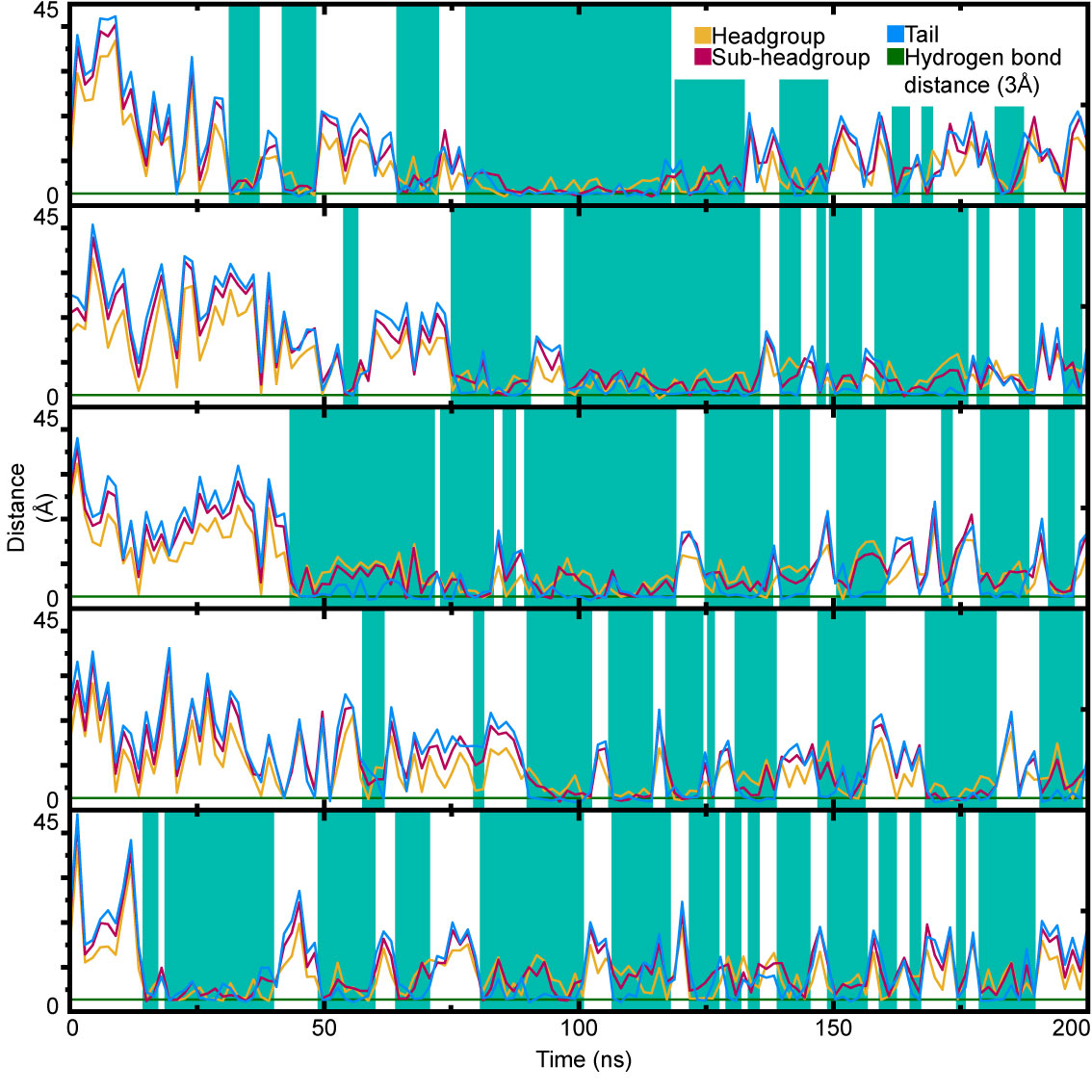

**Figure 8.** A representation of the SaCM system exposed to the CHX in 20% PROH solution at (left) 0, (top right) 100 and (bottom right) 200 ns. Showing head group phosphorous atoms in red, phosphate oxygen in violet and CHX molecules in green

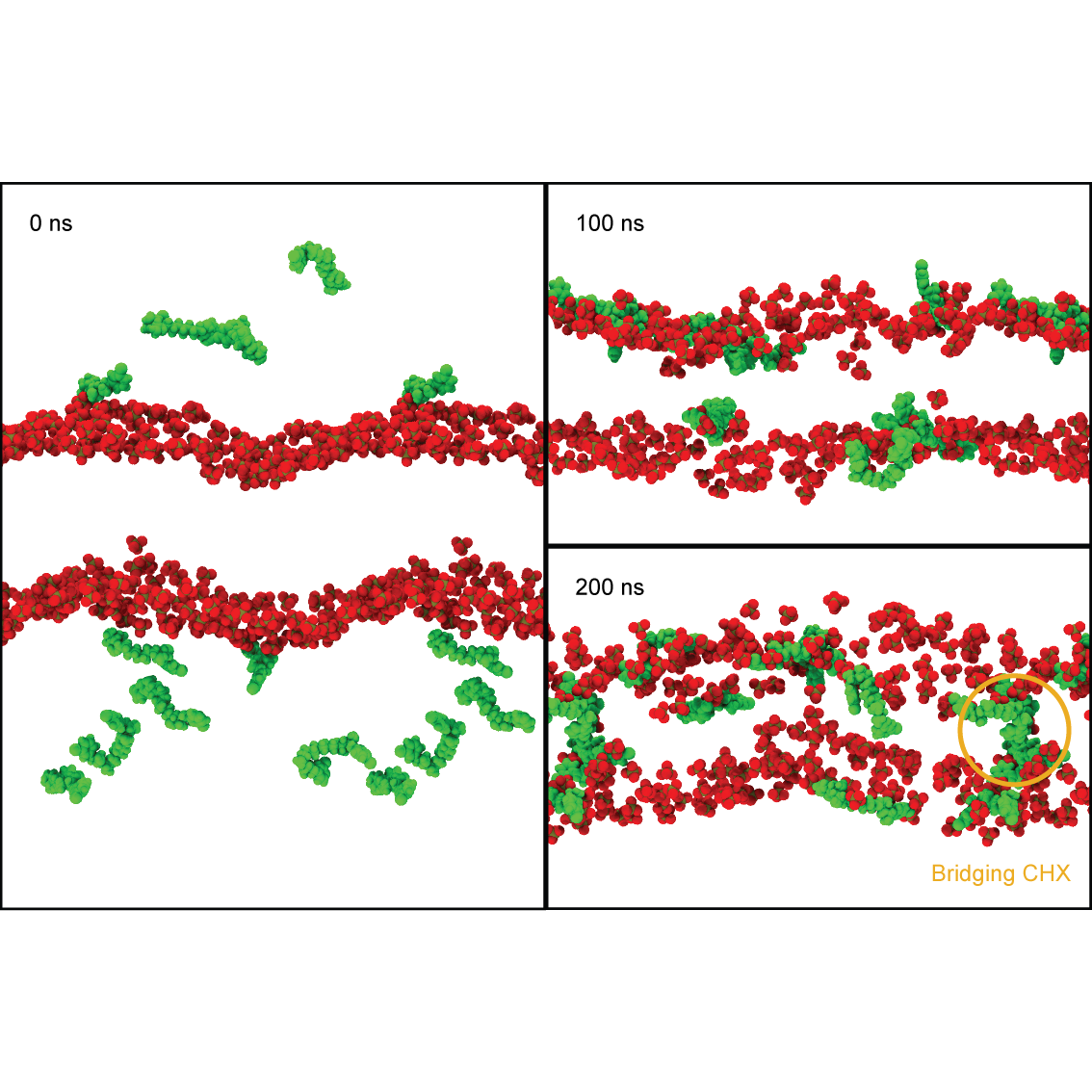

**Figure 9.** The SASA over time of SaCM model lipid membrane systems containing (yellow) PROH and (red) ISOP with CHX in aqueous 0.15 M KCl over 200 ns of production. Visual snapshots of a CHX surface cluster forming in the ISOP system at (green) 20, (cyan) 23 and (pink) 35 ns show carbon CHX residues coloured by residue number, lipid phosphate oxygen in red and phosphate phosphorous in gold. The top row of snapshots look down on the membrane surface along the membrane normal while the lower row look perpendicular to the bilayer normal.

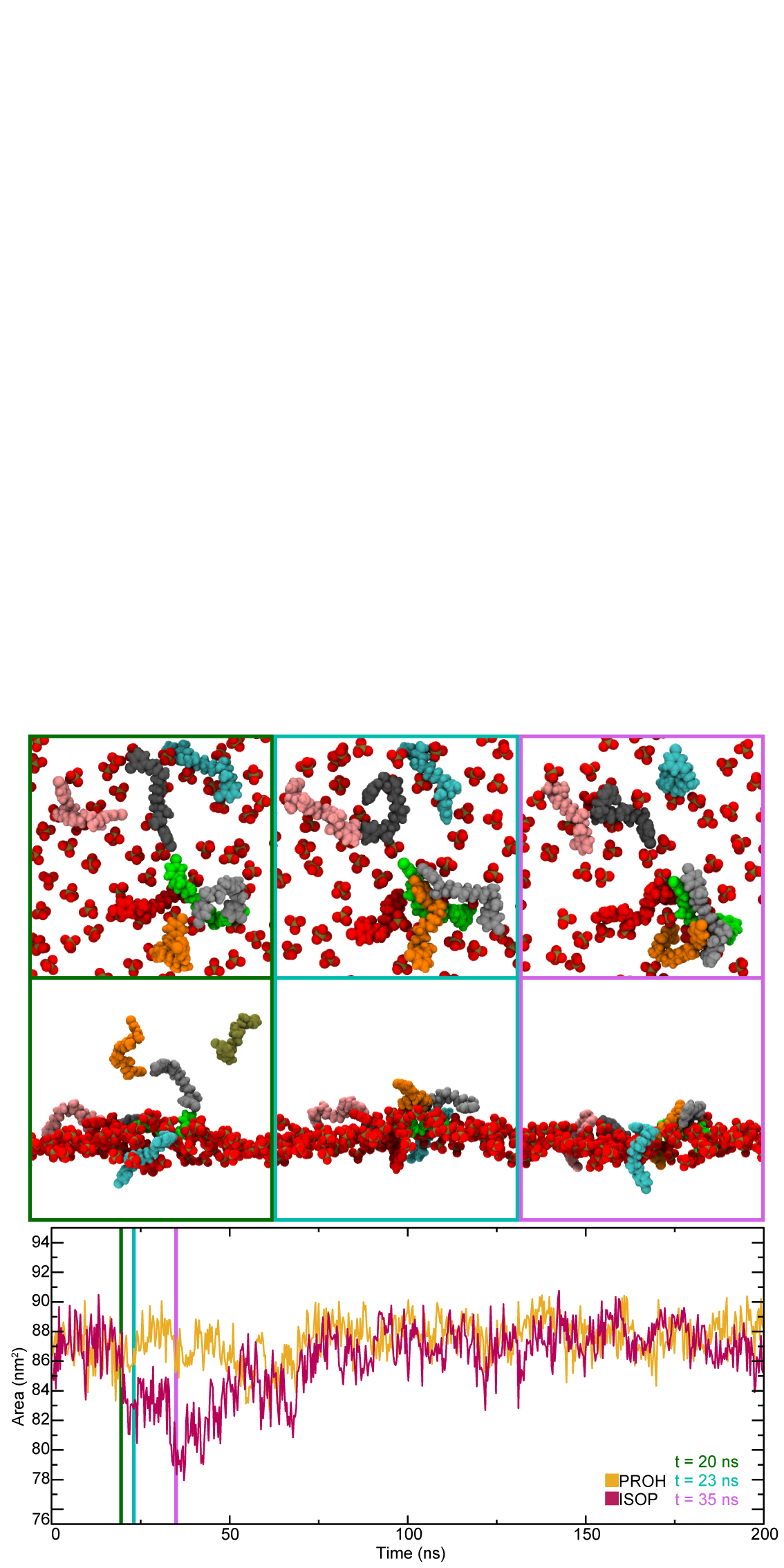

**Figure 10.** A representation of the SaCM after (left) 0, (top right) 100 and (bottom right) 200 ns of exposure to the PROH-CHX solution. Showing CHX in green, lipids as translucent grey, phosphate oxygen in red, phosphate phosphorus in gold and PROH in yellow.

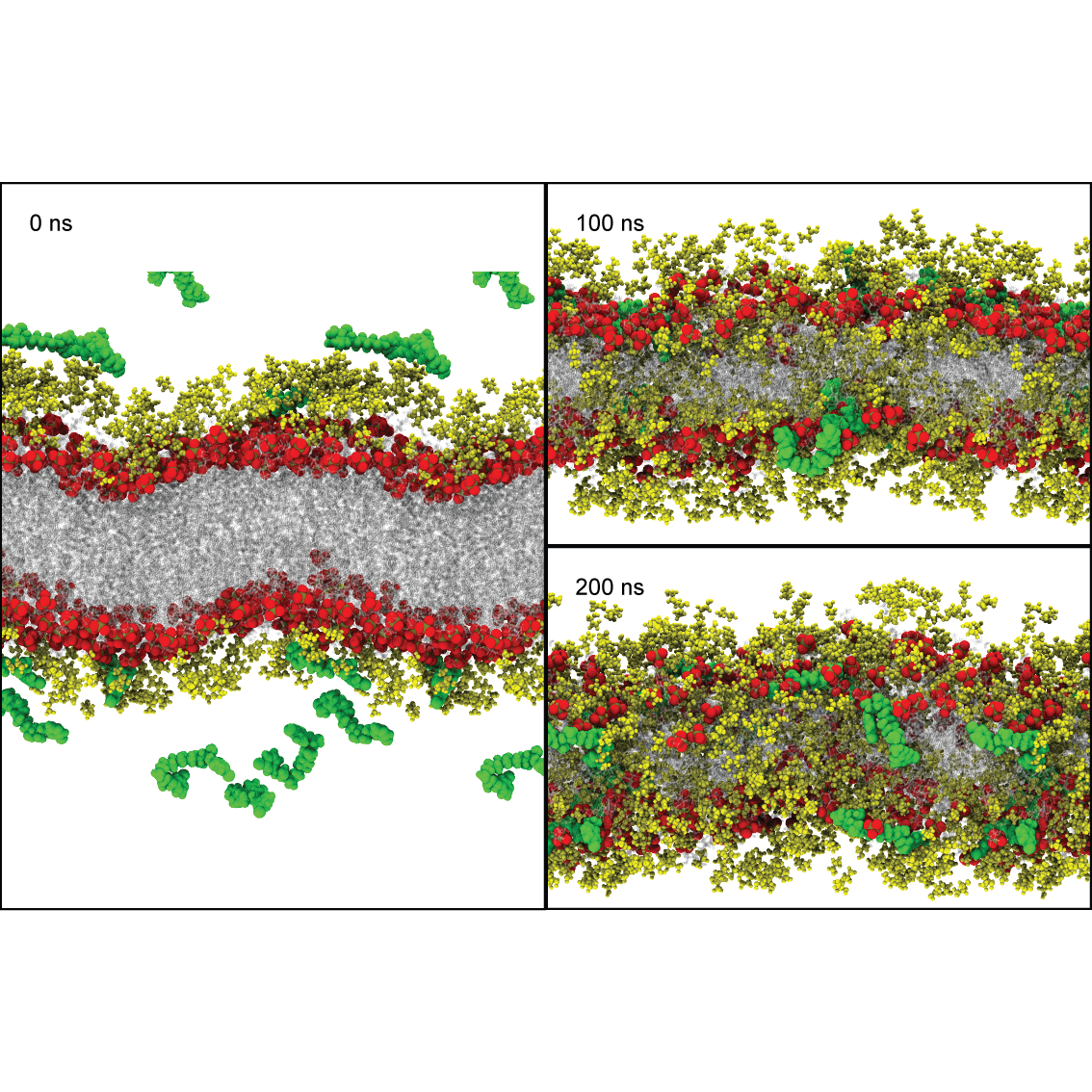

**Figure 11.** A representation of the EcIM after (left) 0, (top right) 100 and (bottom right) 200 ns of exposure to the PROH-CHX solution in a 200 ns production run. Showing CHX in green, lipids as translucent grey, phosphate oxygen in red, phosphate phosphorous in gold and PROH in yellow.

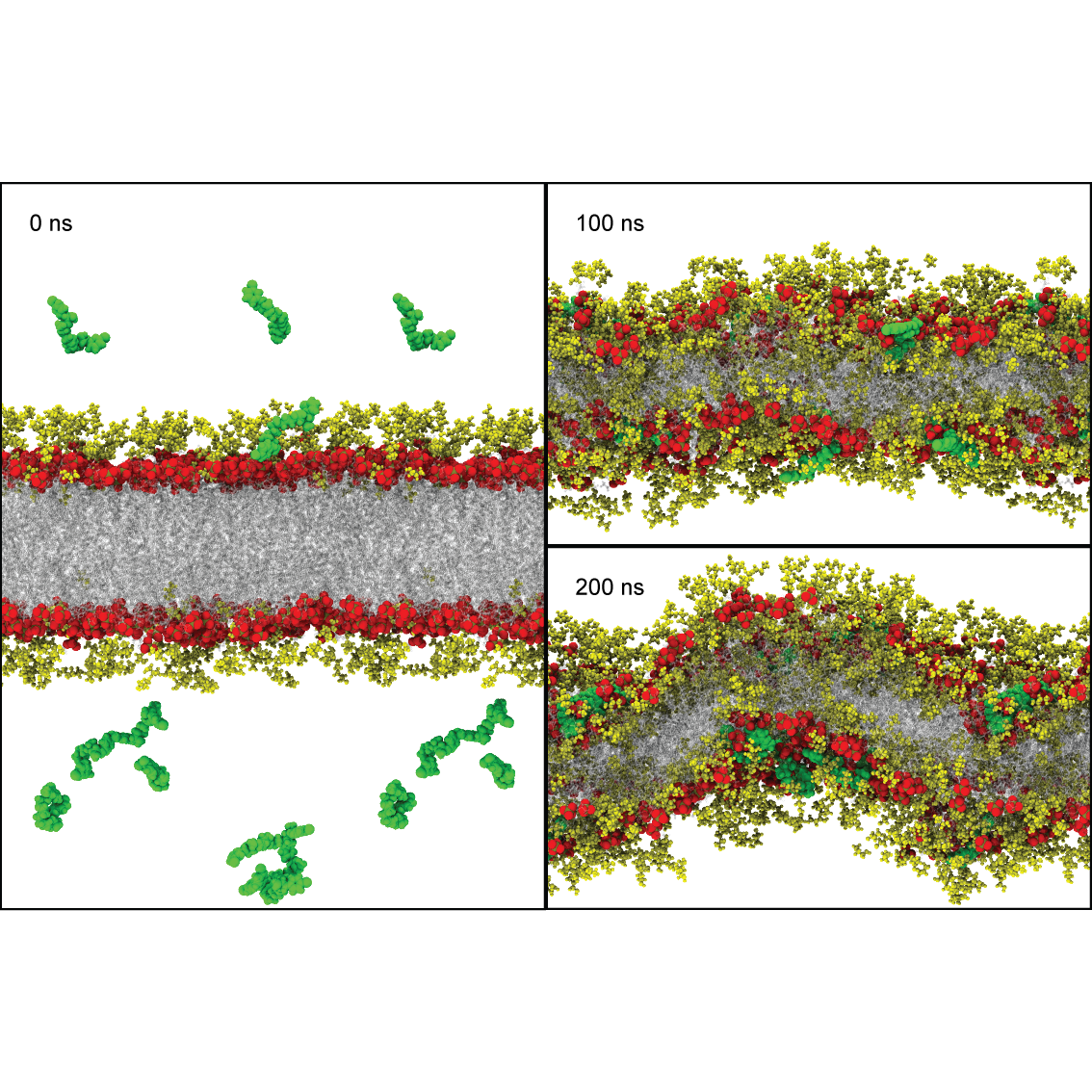

**Figure 12.** Contact mapping of (A) PE, (B) PG and (C) DPG in the EcIM bilayer between 100 and 200 ns with (1) PROH and (2) water mapped linearly from (blue) o contact to (red) the most seen by any atom in the lipid.

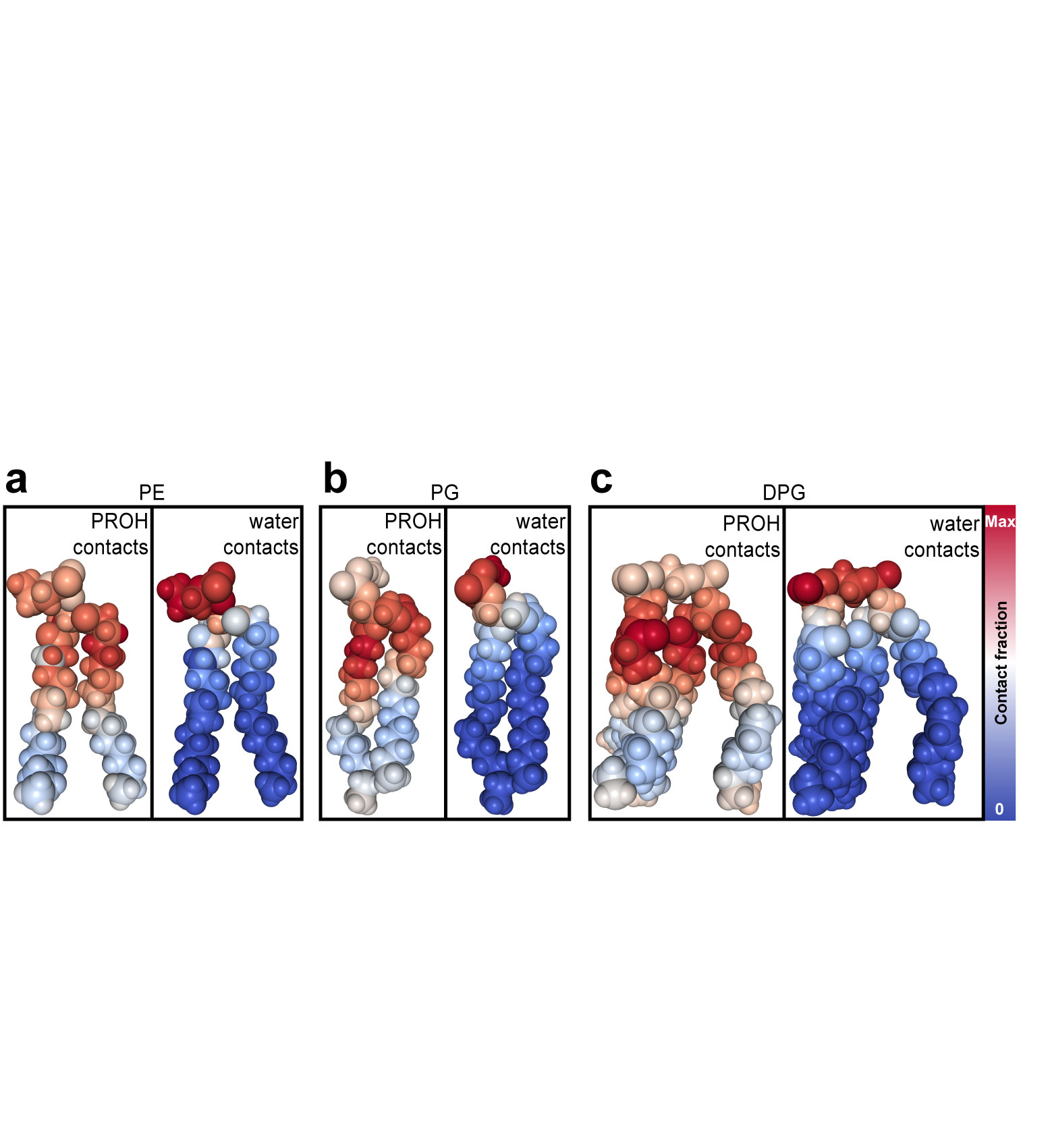

**Figure 13.** Density analysis along the z-axis of the EcIM system exposed to (A) PROH and (B) ISOP solutions. Analysed at time intervals (1) 0-20 and (2) 80-100 ns during a 200ns production run.

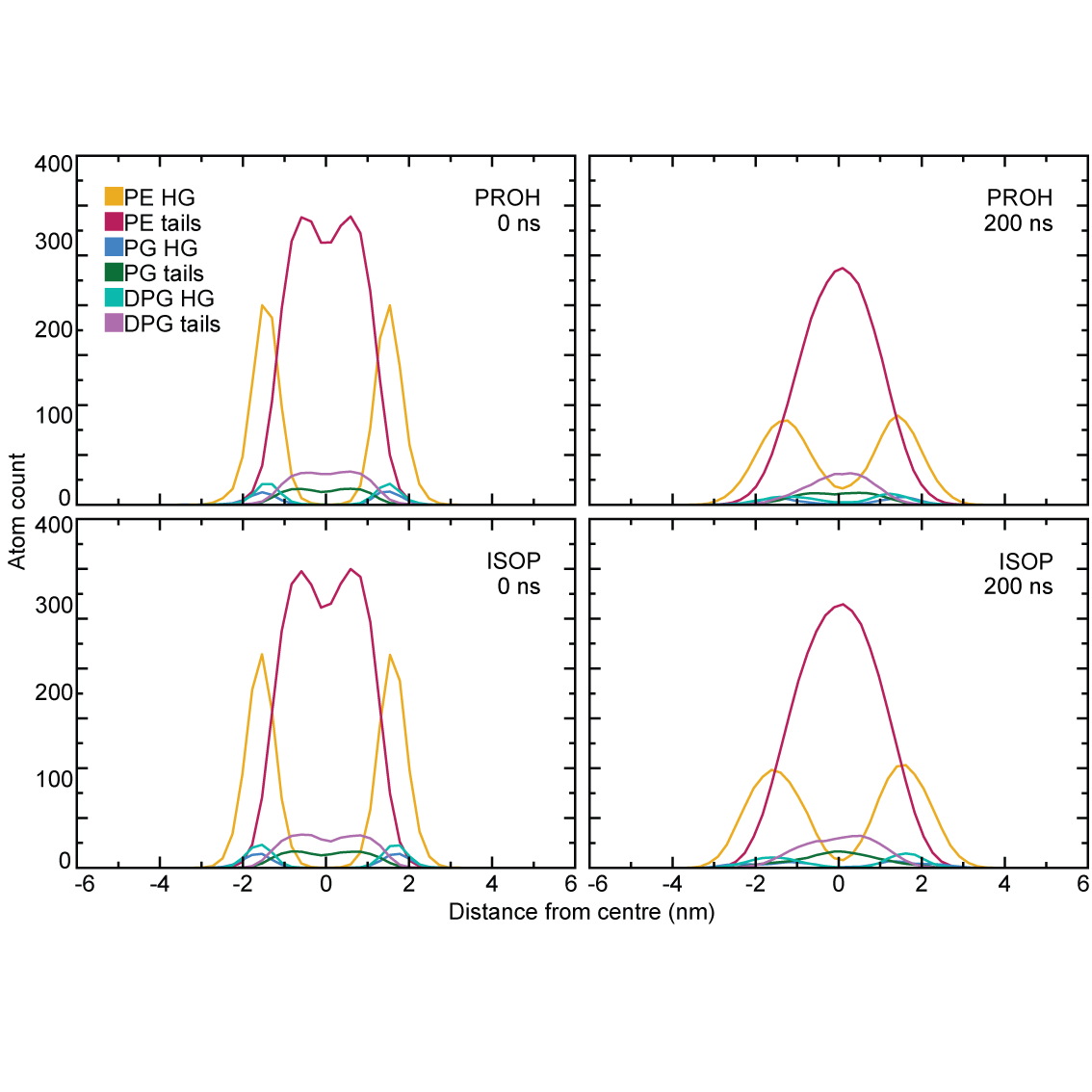

**Figure 14.** Density analysis along the z-axis of the EcIM systems showing specifically alcohol and CHX when the EcIM was exposed to PROH and ISOP solutions with average head group position indicated by dotted lines.

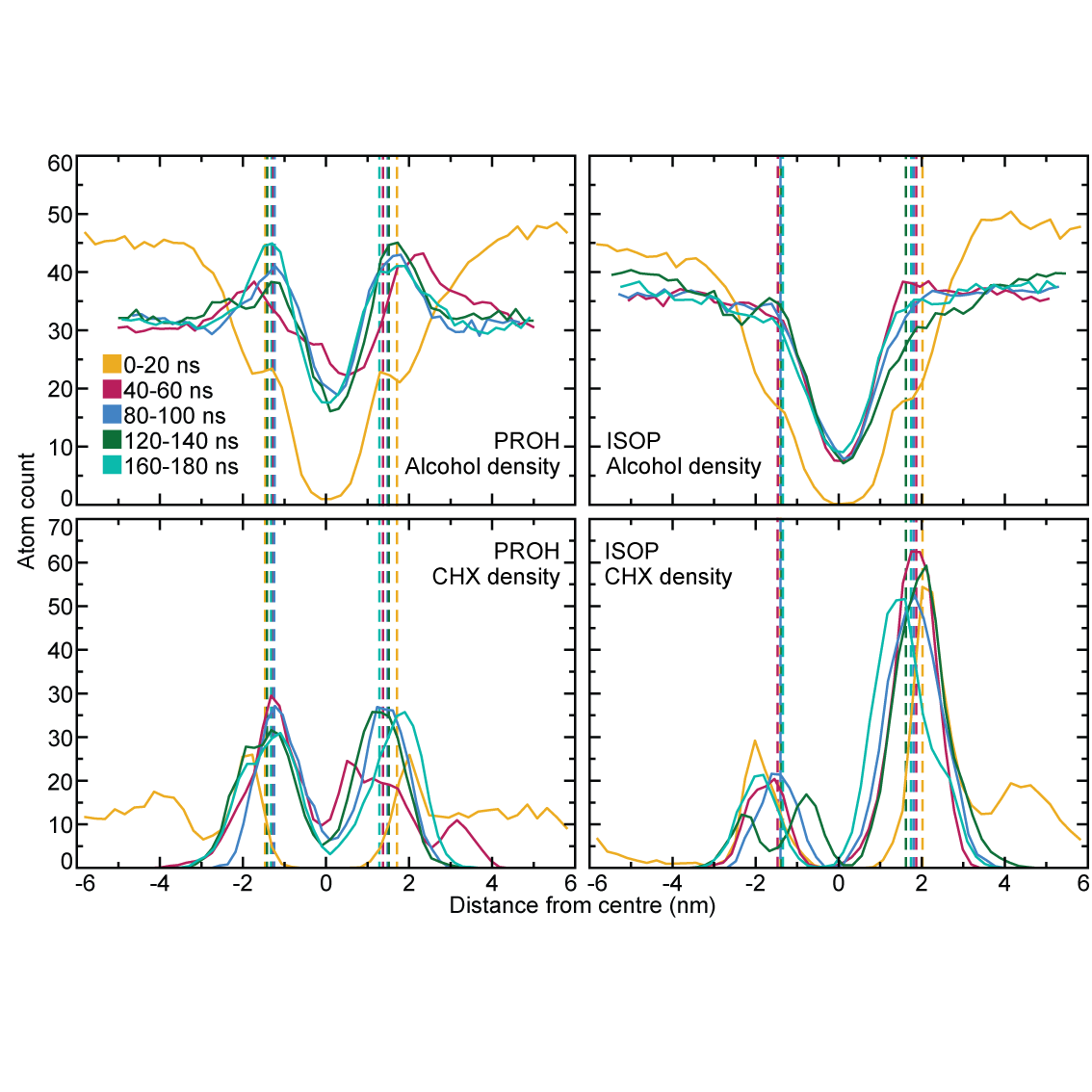

**Figure 15.** The SASA of all CHX combined in the EcIM systems containing 20% (yellow) PROH and (red) ISOP over the whole 200 ns of production.

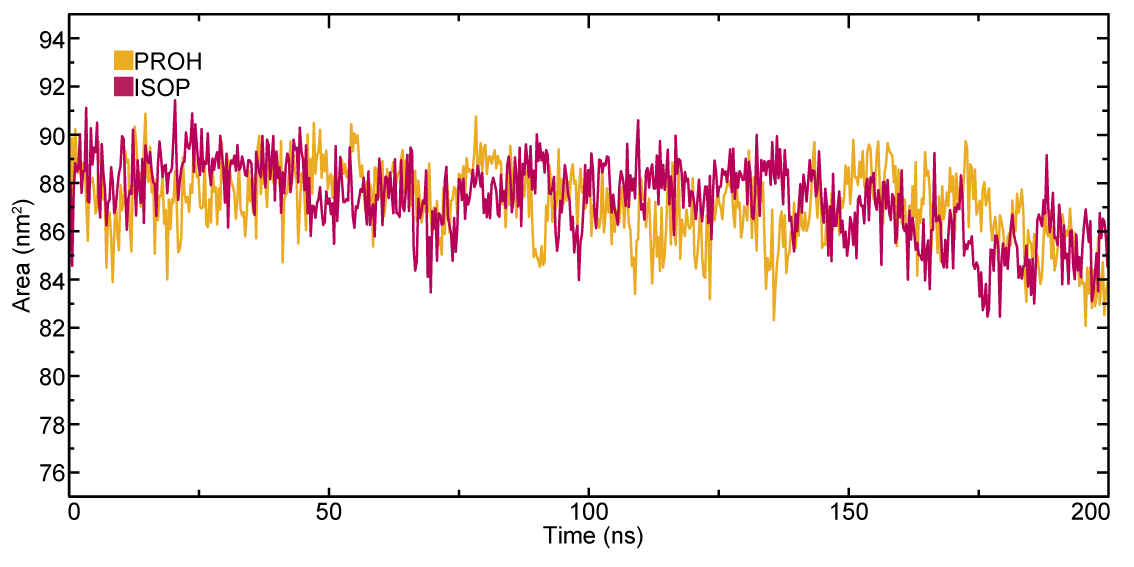

**Figure 16.** Snapshots of the CHX molecule placed in the membrane centre of a pore induced by a 0.15 V nm^-1^ field which was turned off at t = 0 ns at (top left) 0, (top centre) 10, (top right) 20, (bottom left) 30, (bottom centre) 40 and (bottom right) 50 ns. Showing phosphorous in gold, oxygen in red, carbon in cyan, hydrogen in white, nitrogen in blue and lipids as translucent grey.

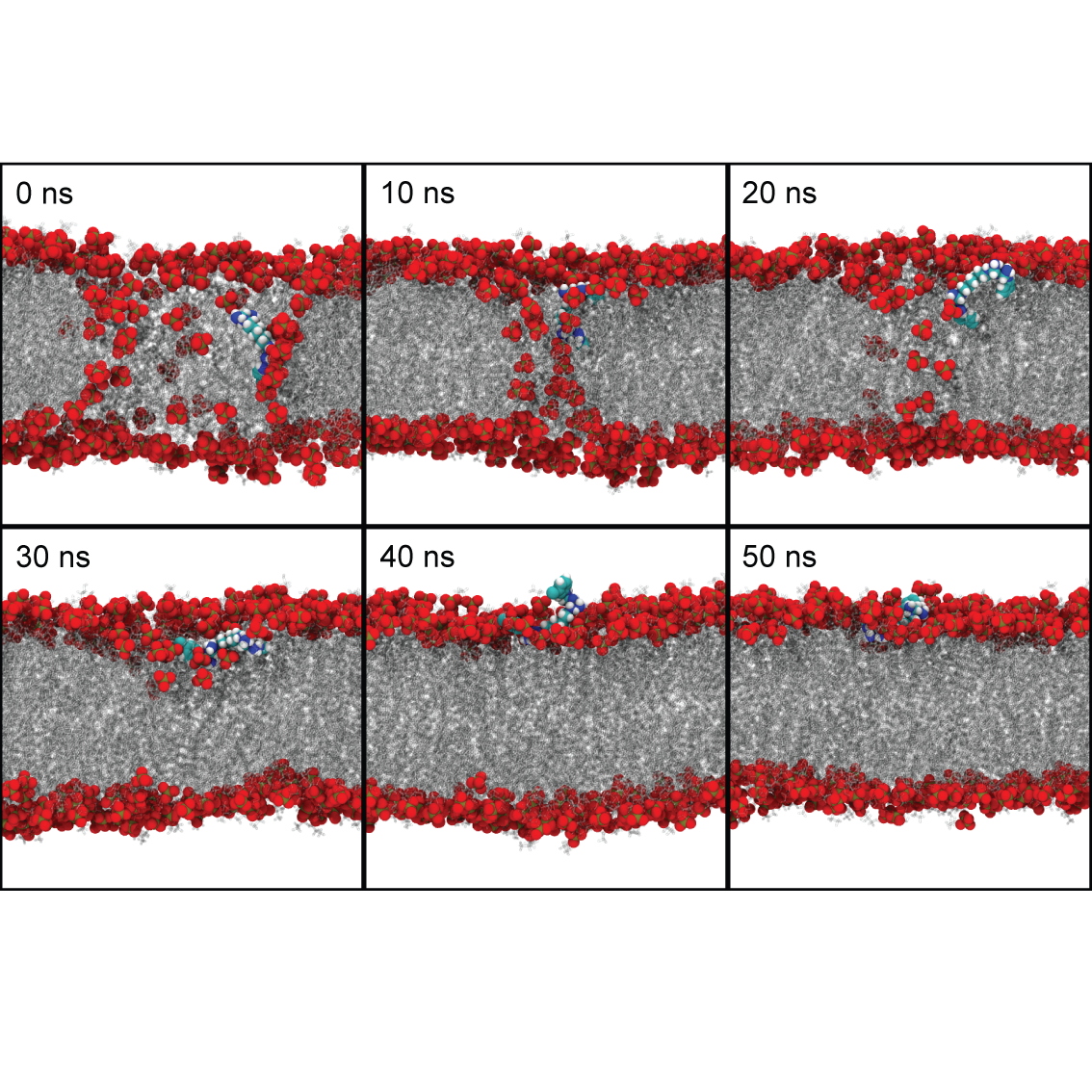

**Figure 17.** Snapshots of of GLUC interacting with the EcIM, showing (A) the membrane phosphates as well as GLUC at 100 ns, (B) PVCL2 interacting with GLUC at the headgroup and (A) PE interacting with GLUC at the headgroup. Phosphorous in gold, oxygen in red, carbon in cyan, hydrogen in white, nitrogen in blue (in A, GLUC is pink, whereas in B and C, GLUC is highlighted by yellow circles).

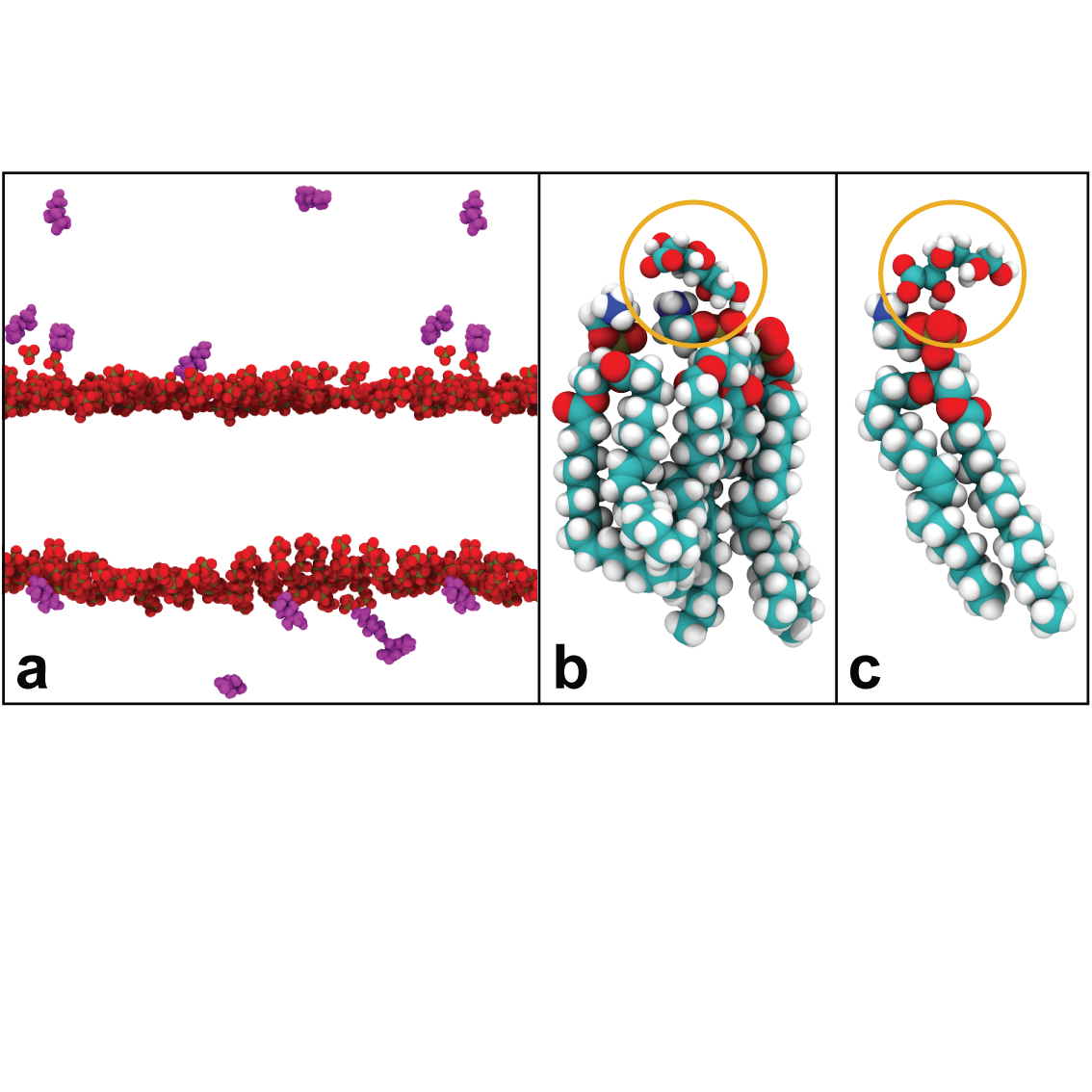

**Figure 18.** The minimum distance between 9 seperate GLUC and all CHX molecules in an MD system containing 20 GLUC and 10 CHX in 0.15 M KCl with an EcIM model.

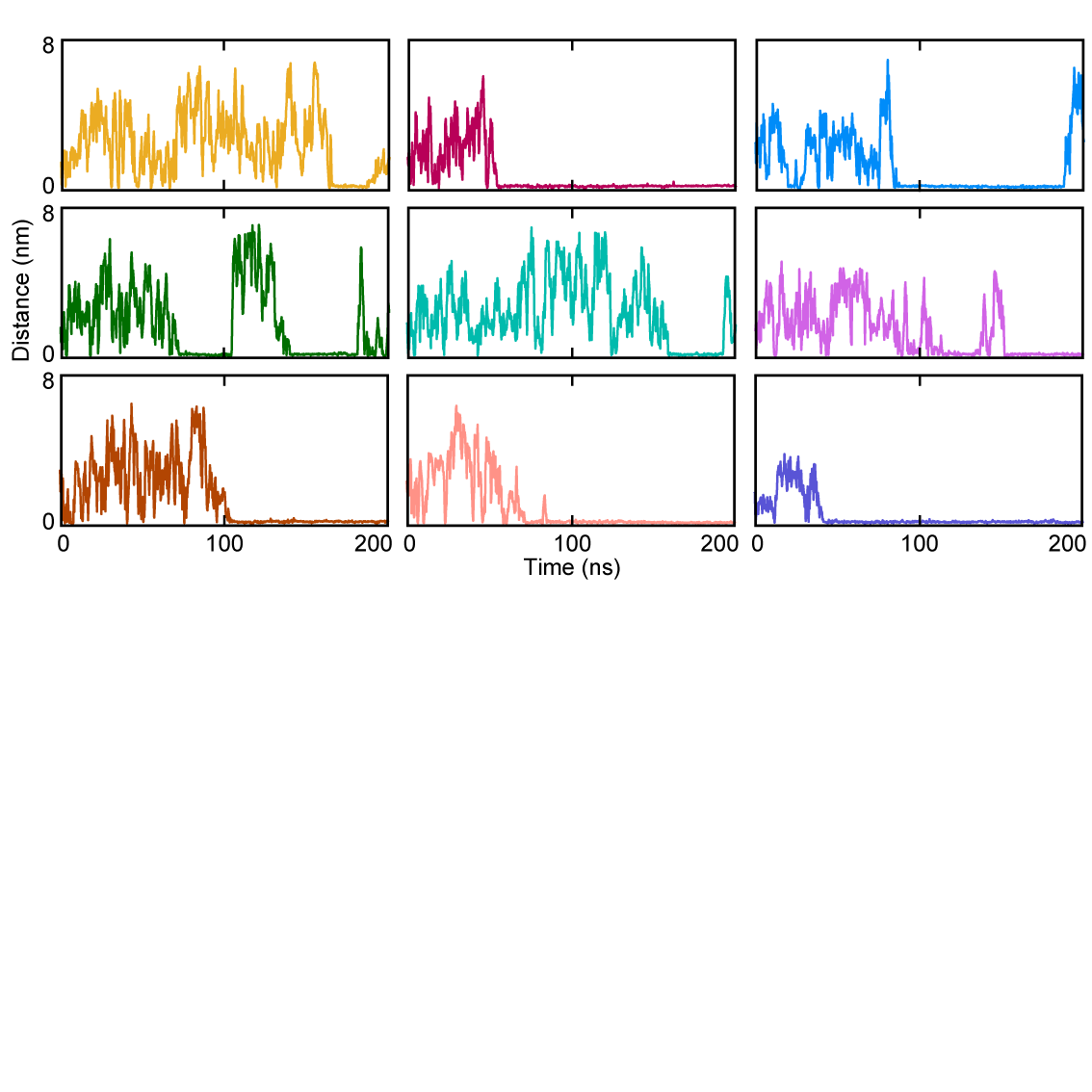

**Figure 19.** Density analysis along the z-axis of the EcOM system exposed to PROH and ISOP solutions. Analysed at time intervals 0-20 and 180-200 ns during a 200ns production run, showing the density shoulder caused by the ‘budding’ effect.

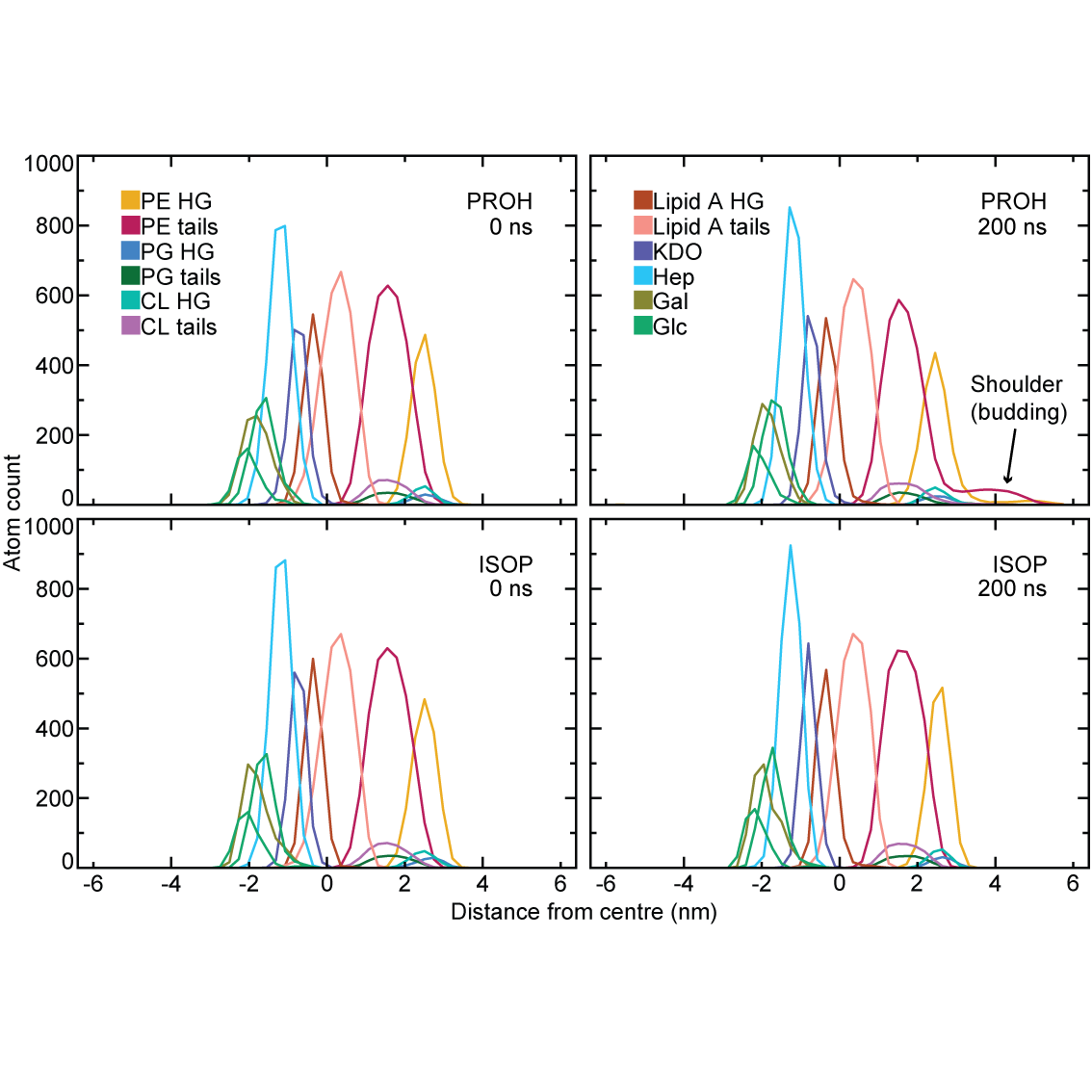

**Figure 20.** The SASA over time of all CHX molecules combined present in the EcOM membrane systems exposed to 20% (yellow) PROH and (red) ISOP over 200 ns of production.

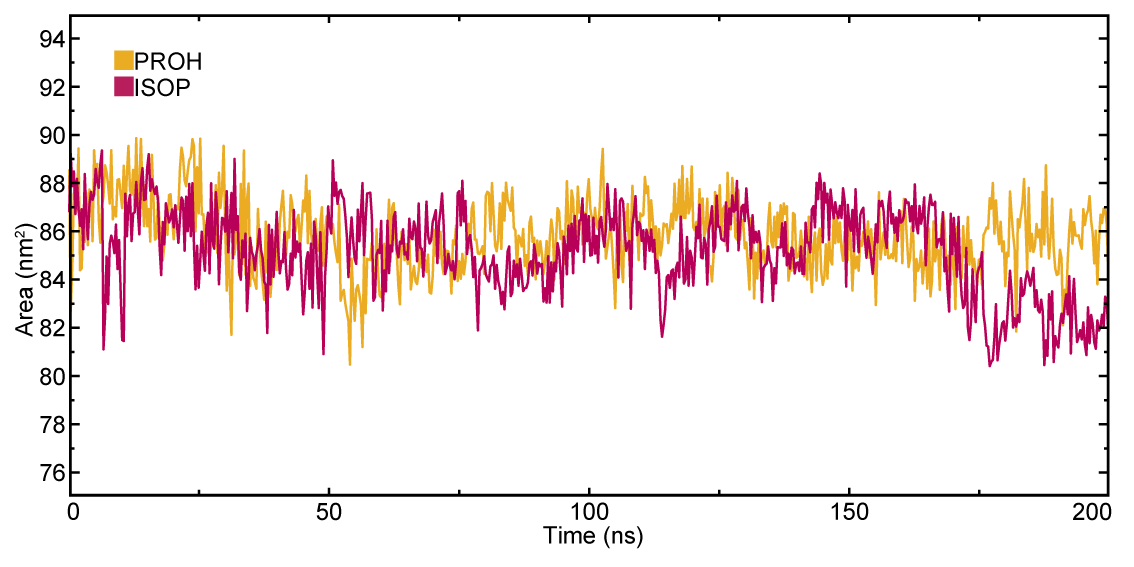

**Figure 21.** The tracked (a) time and (b) z-axis coordinate of a CHX molecule which was seen to interact with the LPS leaflet for an extended period, with the x and y coordinate of CHX recorded, corresponding to the lateral dimensions of the membrane while an increasing z-coordinate corresponds to CHX moving into the LPS leaflet which was facing downwards in the simulation box.

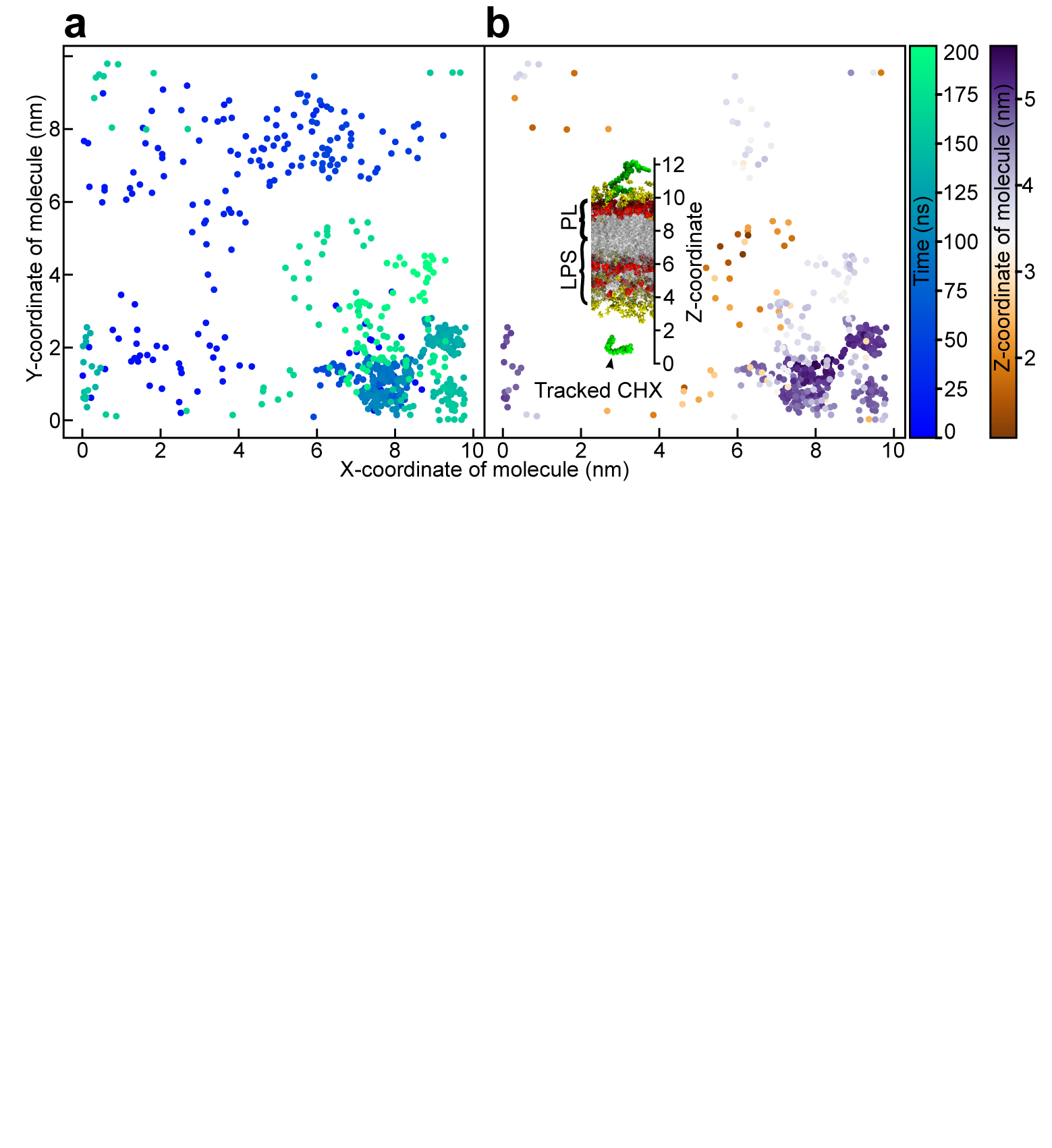

**Figure 22.** Snapshots of the Ra-LPS leaflet of the EcOM after (a) 100 and (b) 200 ns of exposure to the solution of 20% PROH and CHX, with snapshots of the CHX-sugar interactions at (c) 80, (d) 100, (e) 120 and (f) 140 ns in a 200 ns production run. Showing carbon in cyan, hydrogen in white, nitrogen in blue, chlorine in pink, lipid-A as translucent, ketodeoxyoctonic acid in dark blue, heptose in purple, glucose in green and brown and galactose in bright pink.

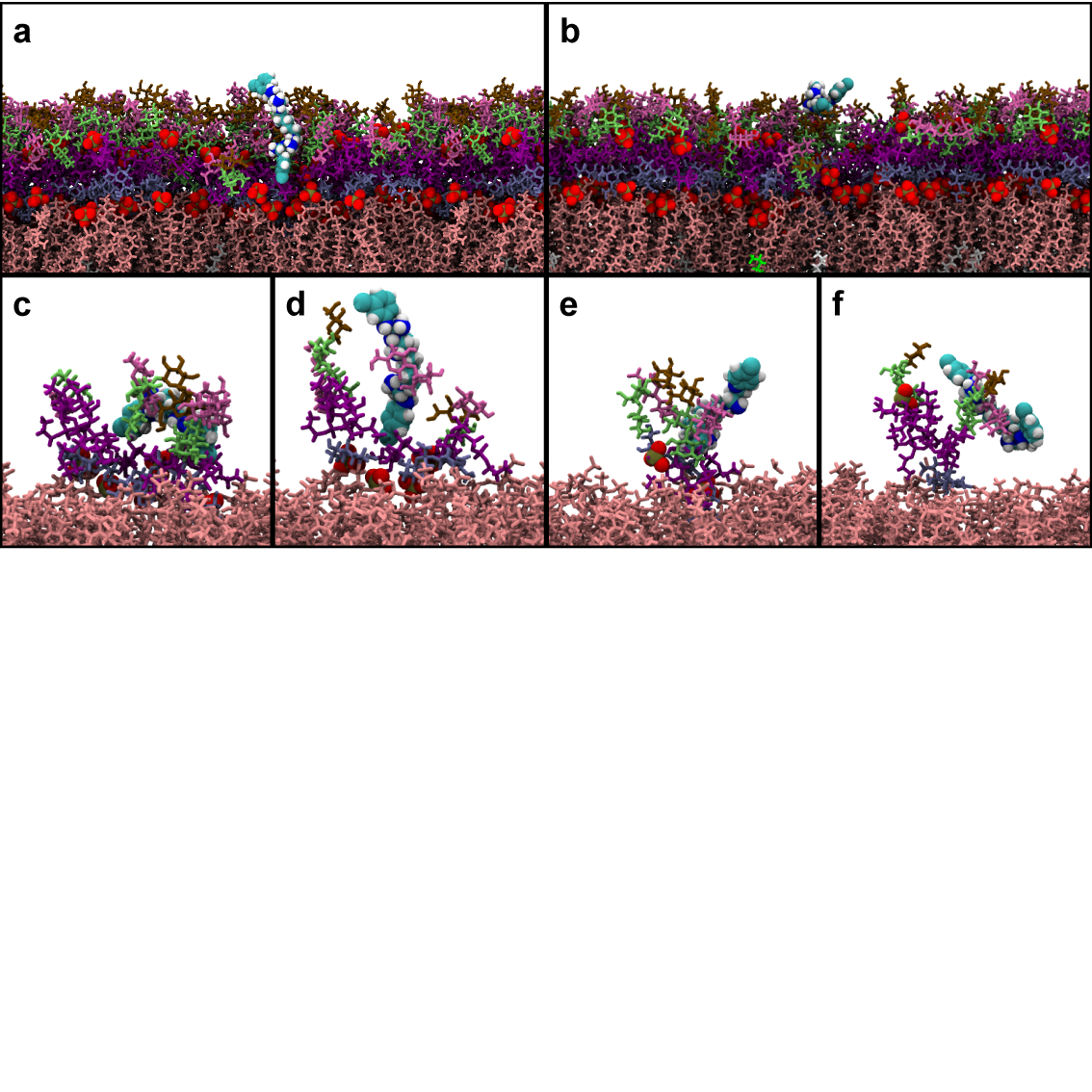

**Figure 23.** (a) The average RDF of each CHX molecule in the system with respect to all other CHX molecules in the system averaged over the final 20 ns of the 200 ns production run with a cut-off of 2.4 nm. (b) The total SASA of all CHX molecules in the system over the full 200 ns extracted trajectory of just CHX. (c) Snapshots of the 1 M KCl system at 0ns, 63 ns and 145 ns representing points at which SASA was particularly low.

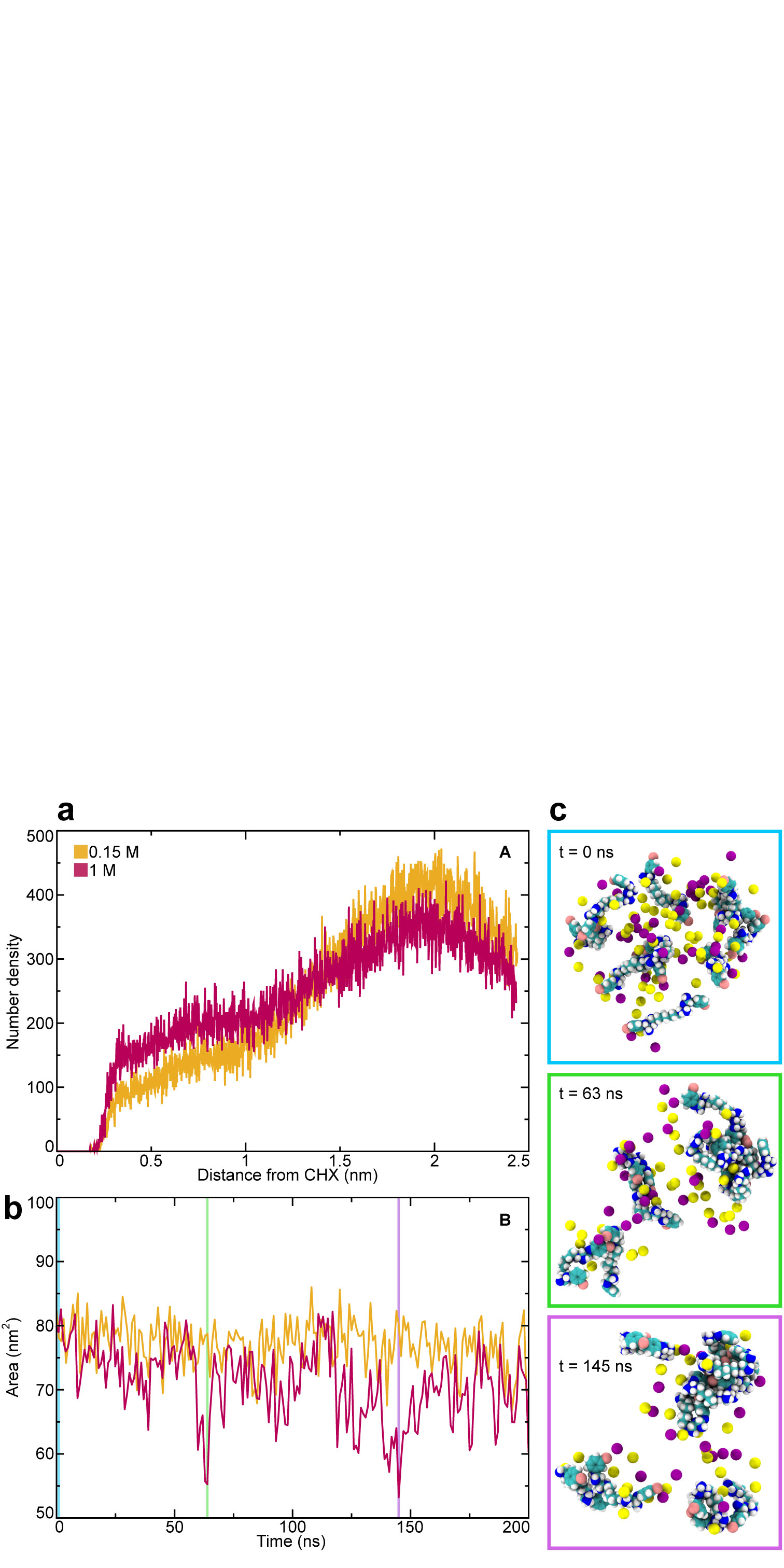

**Figure 24.** Cluster analysis showing the presence of the 5 most common clusters of CHX overlayed onto the RMSD of all conformations detected to show how common they are in the full production run in the (a) 0.15 and (b) 1 M KCl systems.

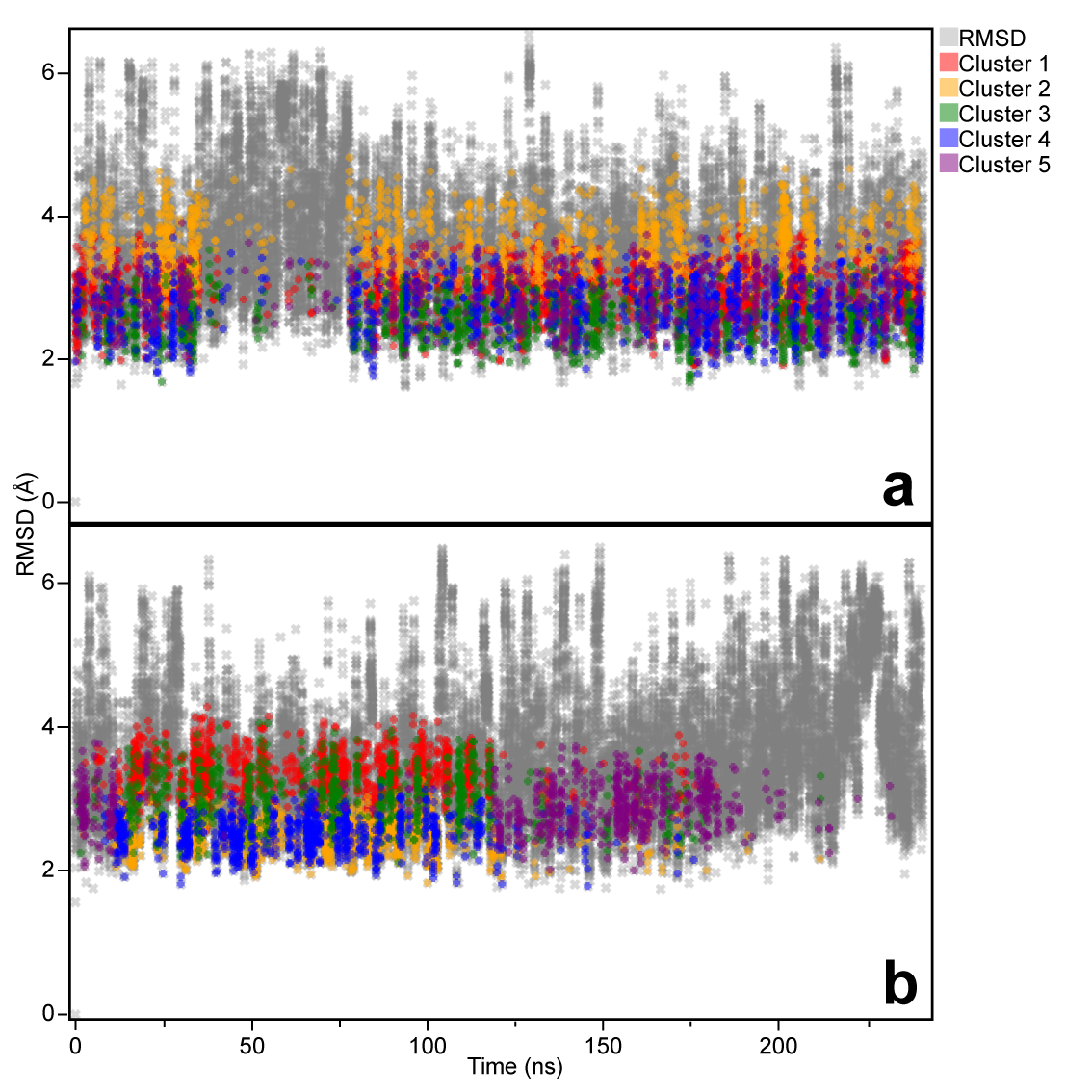

$$\boldsymbol{a}\left( \boldsymbol{\tau}_{\boldsymbol{m}} \right)\boldsymbol{=A}\left( \boldsymbol{\tau}_{\boldsymbol{m}} \right)\boldsymbol{\times}\boldsymbol{A(0)}^{\boldsymbol{-1}}$$

(1)

$$\boldsymbol{a}\left( \tau_{m} \right)=\boldsymbol{X D} \boldsymbol{X}^{-1}$$

(2)

$$\boldsymbol{R}=\frac{\boldsymbol{X} \boldsymbol{D} \boldsymbol{X}^{-1}}{\tau_{m}}$$

(3)

**Equation 1.** Full matrix approach to calculating the magnetisation exchange rate between molecules in a 2D NOESY NMR experiment, where a bold variable denotes a matrix.
